# Molecular fingerprints of cell size sensing and mating type differentiation in pennate diatoms

**DOI:** 10.1101/2024.04.15.589526

**Authors:** Darja Belišová, Gust Bilcke, Sien Audoor, Sofie D’hondt, Lieven De Veylder, Klaas Vandepoele, Wim Vyverman

## Abstract

- A unique cell size sensing mechanism is at the heart of the life cycle of diatoms. During population growth, cell size decreases until a Sexual Size Threshold (SST) is reached, below which cells become sexually competent. In most pennate diatoms, the two mating types undergo biochemical and behavioral differentiation below the SST, although the molecular pathways underlying their size-dependent maturation remain unknown.
- Here, we developed a method to shorten the generation time of *Cylindrotheca closterium* through single-cell microsurgery, enabling the transcriptomic comparison of genetically identical large and undifferentiated cells with small, sexually competent cells for six different genotypes.
- We identified 21 genes upregulated in small cells regardless of their mating type, revealing how cells undergo specific transcriptional reprogramming when passing the SST. Furthermore, we revealed a size-regulated gene cluster with three mating type-specific genes susceptible to sex-inducing pheromones. In addition, comparative transcriptomics confirmed the shared mating type specificity of Mating-type Related Minus 2 homologs in three pennate diatoms, suggesting them to be part of a conserved partner-recognition mechanism.
- This study sheds light on how diatoms acquire sexual competence in a strictly size-dependent manner, revealing a complex machinery underlying size-dependent maturation, mating behavior, and heterothally in pennate diatoms.

## Introduction

Diatoms are an extremely successful group of unicellular primary producers in freshwater and marine environments (Field *et al*., 1998). Despite their extraordinary species diversity underlying the colonization of widely different habitats spanning terrestrial to aquatic realms, all diatoms share a unique silica cell wall consisting of two halves (thecae) slightly differing in size. During cell division, the two daughter cells each inherit one of the rigid parental thecae that are dissimilar in size and fit as a “lid on a box” (**Figure 1**). As a result, an asexually growing population of cells will gradually decrease in mean cell size, a phenomenon known as the MacDonald-Pfitzer rule (Pfitzer, 1869; Macdonald, 1869). Different mechanisms of cell size restoration mechanisms have been described (Nagai *et al*., 1995; Chepurnov *et al*., 2004; Kaczmarska *et al*., 2022); but most commonly this happens via the sexual phase during which diploid vegetative cells form haploid gametes. The ability to engage in sexual reproduction is strictly cell-size controlled and triggered once cells drop below a species-specific Sexual Size Threshold (SST). The cellular mechanisms and molecular changes associated with this unique ability of diatom cells to sexually mature by measuring their own cell size are still completely unknown.

**Figure 1:**
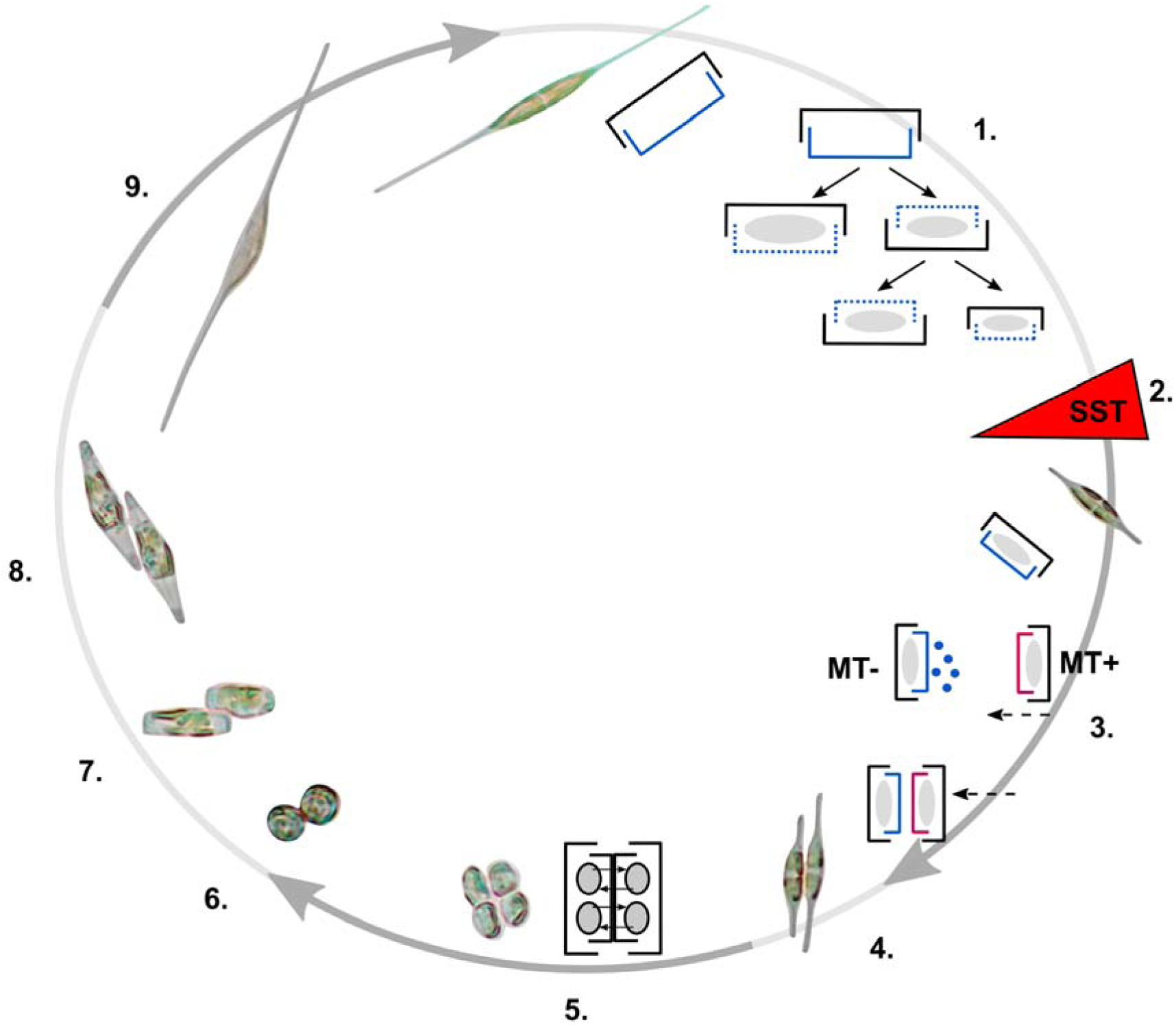
**A typical life cycle of a raphid pennate diatom**, illustrated with the species *C. closterium*. (**1**) Cells divide vegetatively and the mean size in the population drops below the sexual size threshold (SST) (**2**). Small cells become sexually mature and seek a compatible mate (**3**). In *C. closterium*, mating type minus (MT-) produces pheromone (SIP-, blue dots) while mating type plus (MT+) evinces oriented motility towards the partner (**3**,**4**). (**5**) Each gametangial cell produces two gametes. (**6**) Gametes fuse and form zygotes (**7**). Zygotes expand and become auxospores (**8**), and finally, an initial cell (**9**) starts a new generation.

In the generally homothallic centric diatoms, cells produce oogonia and sperm cells, while in raphid pennate diatoms vegetative cells become responsive to sexual cues after passing the SST and transform into morphologically identical but functionally differentiated gametangial cells (Moeys *et al*., 2016; Basu *et al*., 2017; Bilcke *et al*., 2021; Klapper *et al*., 2021). Recent advances revealed the complex nature of diatom sexual behavior, including the identification of multiple sex pheromones involved in the mating process (Gillard *et al*., 2013; Moeys *et al*., 2016; Klapper *et al*., 2021, 2023).

Most raphid pennate diatoms have a heterothallic mating system with two mating types (MT+and MT-) (reviewed in (Chepurnov *et al*., 2004; Davidovich *et al*., 2015; Poulíčková *et al*., 2019; Bilcke *et al*., 2022)). Although the elucidation of the diploid sex-determination system in heterothallic pennate diatoms is still in its infancy, crossing experiments have revealed that one of the two mating types is typically heterogametic, i.e. produces different gametes that determine the mating type of the offspring (Davidovich 2002, Vanstechelman *et al*., 2013). Additionally, recent studies in *Seminavis robusta* and *Pseudo-nitzschia multistriata* have demonstrated the existence of a single sex-determining locus that drives the expression of downstream mating type related genes (Vanstechelman *et al*., 2013; Russo *et al*., 2018). Specifically, the sex-determining A allele of the *P. multistriata* Mating-type Related Plus 3 (*MRP3*) gene segregates with MT+ strains (Russo *et al*., 2018). A simple control mechanism, where differential methylation of the sex locus leads to mono-allelic expression of *MRP3* only in MT+, has been described by Ruggiero et al. (2023). MRP3 acts as a master regulator, activating downstream MT+ genes *MRP1* and *MRP2*, while inactivating the MT-specific genes *MRM1* and *MRM2*. Importantly, despite the widespread occurrence of heterothally in pennate diatoms, no homologs of this sex determining system have yet been identified in other pennate species outside the *Pseudo-nitzschia - Fragilariopsis* clade (Russo *et al*., 2018).

Here, we studied size sensing and mating type differentiation in the raphid pennate diatom *Cylindrotheca closterium,* a common and globally widespread member of marine, brackish water phytoplankton and microphytobenthos (Audoor *et al*., 2024). We developed a protocol to significantly shorten generation time by mechanically reducing cell length, allowing us to compare gene expression of cells above and below the SST for the same genotypes. We carried out differential gene expression analysis using robust transcriptome profiling of six different genotypes, encompassing both mating types in both size variants. Phylogenetic analysis and cross-species transcriptomic analysis were used to reveal conserved size- and mating type-specific genes.

## Materials & Methods

### Culture conditions for *C. closterium* and *S. robusta* cells

Cultures of *C. closterium* and *S. robusta* were grown in artificial seawater (ASW) composed of Tropic Marin BIO-ACTIF sea salt (34.5 g/l; Tropic Marin, Wartenberg, Germany) and sodium bicarbonate (0.08 g/l) supplemented with Guillard’s F/2 Marine Water enrichment solution (Sigma-Aldrich, Belgium) and an antibiotic mixture (500 mg/l penicillin, 500 mg/l ampicillin, 100 mg/l streptomycin and 50 mg/l gentamicin). Cells were sustained under constant illumination (25 µmol/m2/s) at 18°C, if not stated otherwise. An initial set of strains was provided by the Belgian Coordinated Collection of Microorganisms (BCCM/DCG) (https://bccm.belspo.be/about-us/bccm-dcg) (**Supplementary Table 1**).

### Experimental size manipulation and recovery of *C. closterium*

Four parental *C. closterium* strains of two different mating types were crossed to create progeny strains. More information about the parental strains can be found in **Supplementary Methods**. Single cells were isolated to start monoclonal progeny strains and subsequently their subpopulations were reduced in size using a novel microsurgery approach. Particularly, long apices (cell protrusions) of single cells were cut off using an extremely thin tip of a glass Pasteur pipette, with an open diameter ranging from 50 to 200 µm (**Supplementary Figure 1**). Prior to the experiment, pipettes used for cutting and isolation were manually thinned using a Bunsen Burner and clipped by a tweezer to create open and sharp tips (**Supplementary Video**). In healthy and unstressed cultures, ∼50-70% of cells recovered successfully from the cutting procedure, forming new monoclonal, smaller-celled populations. Such manipulated single cells were immediately isolated in a 96 well-plate (Greiner CELLSTAR®, Sigma Aldrich Belgium) and were then maintained two weeks in low light conditions (10 – 15 µmol photons/m2/s) for wounding recovery, giving rise to new monoclonal populations that were genetically identical to their larger counterparts but with a smaller cell size. After this immediate recovery period, cultures were maintained in the growth phase for at least three months to ensure their complete recovery. An identical period of vegetative growth was enabled for large non-manipulated paired cultures. During this extended period of growth, cultures were grown in 24-well plates under 12/12 day/night cycles. Once a week, each culture was scraped from the bottom of the well and 15 µl of a dispersed culture was inoculated into 1 ml of fresh medium to ensure that cultures remained in exponential growth phase. Finally, all cultures were transferred to small culture flasks (25 cm^2^, VWR ®) to be grown under continuous light for at least four weeks to acclimate before harvesting for the RNA-seq experiment.

### Determination of mating types and selection of genotypes for RNA-sequencing

Before selecting *C. closterium* cultures for the RNA-seq experiment, the average length of cells (the distance between the tip of each apex, **Supplementary Figure 2**) was determined and the mating type was identified by backwards crossing of each culture with both parental genotypes below the SST (Audoor *et al*., 2024). Specifically, synchronized cultures were dark arrested for 36 hours followed by mixing various combinations of two strains in a 96 well-plate with a cell density of 40 cells/µl per strain in a total volume of 200 µl. The potential mating of strains was microscopically inspected after 12, 24 and 48 hours. The experimentally determined mating types are indicated in **Supplementary Table 1**. Backwards crosses were performed for both the small (after microsurgery) and large (original) variants to verify whether small cells can form all sexual cell stages (mating pairs, auxospores, initial cells), while large cells are incapable of sexual reproduction.

Altogether, six *C. closterium* genotypes were selected for RNA-seq of their matched large and small size classes. We picked three MT+ genotypes: C2 (86 µm large cells vs 34 µm small cells), MC4 (68 µm vs 29 µm) and, MC5 (80 µm vs 26 µm); and three MT-genotypes: – A5 (81 µm vs 47 µm), A6 (67 µm vs 37 µm) and, MC3 (74 µm vs 45 µm) (**Supplementary Table 1**). For *S. robusta*, six strains below the SST, three of each mating type, were selected for additional RNA-seq (**Supplementary Table 1**) to complement existing datasets (Osuna-Cruz *et al*., 2020).

### RNA extraction and sequencing

Exponentially growing cultures of *C. closterium* and *S. robusta* were harvested on membrane filters (3µm, 25mm, Versapor®, Pall Laboratory, VWRTM) and RNA was extracted using a Qiagen RNeasy extraction kit with optimizations. Detailed protocols can be found in **Supplementary Methods**.

Samples with the highest RNA integrity number and sufficient quantity were processed by TruSeq Stranded mRNA Library Prep with polyA selection. The library was further subjected to 2×150bp paired-end sequencing using the Illumina NovaSeq 6000 instrument at VIB Nucleomics Core (Leuven, Belgium www.nucleomics.be). On average, 33.8 ± 4.4 million and 32.0 ± 3.7 million reads per sample were sequenced for *C. closterium* and *S. robusta* respectively.

### Differential expression analysis and functional annotation

Quality trimmed paired-end reads (as explained in **Supplementary Methods**) were mapped to transcripts belonging to the gene annotation model v1.2 of the *C. closterium* reference genome (Audoor *et al*., 2024) using Salmon v1.3 (Patro *et al*., 2017). Read counts were imported into R with tximport v1.22.0 (Soneson *et al*., 2016), followed by filtering, normalization and generalized linear model fitting as detailed in the **Supplementary Methods**. For *C. closterium*, we defined four contrasts to execute a differential expression (DE) analysis with EdgeR v3.36.0 (Robinson *et al*., 2009) (**Figure 2**): (1) cell sizes above and below the SST irrespective of the MT, (2) cells of MT+ versus MT-irrespective of the size, (3) cells of MT+ below the SST versus all other samples, and (4) cells of MT-below the SST versus all other samples. For the “Size” contrast (1), an additional paired analysis was performed, controlling for the variable “genotype” in the generalized linear model. Next, likelihood ratio tests were performed using EdgeR, and p-values were controlled at the 5 % false discovery rate (FDR) level using the Benjamini-Hochberg correction. The analysis of existing (Osuna-Cruz *et al*., 2020) and newly generated RNA-seq data of *S. robusta* used for comparative purposes is explained in **Supplementary Methods**. In addition, differential expression calls during the different stages of sexual reproduction of *C. closterium* and *S. robusta* were retrieved from (Audoor *et al*.,2024).

**Figure 2:**
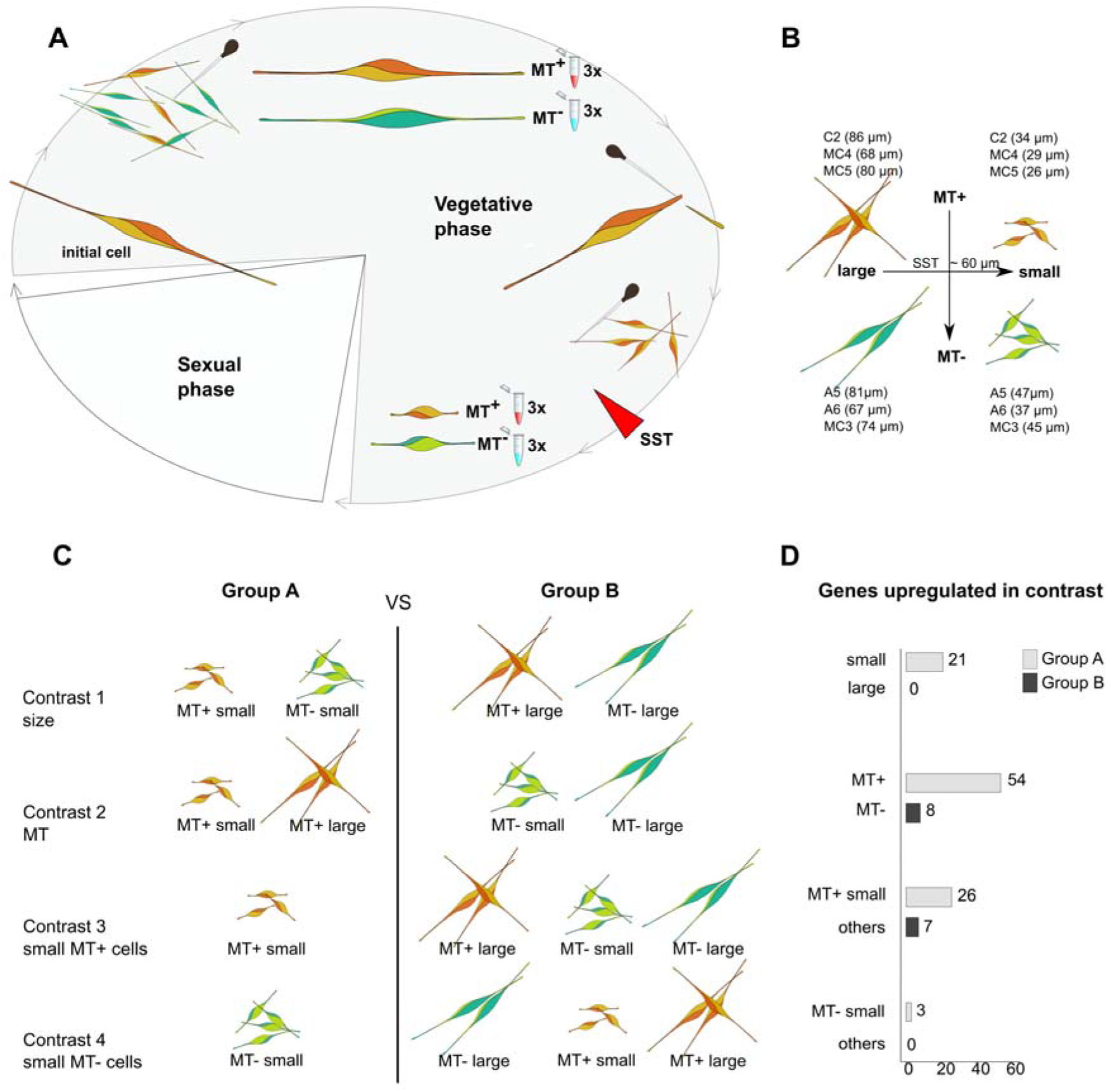
Illustration of the microsurgery-based set-up for size manipulation of *C. closterium*, experimental design of RNA sequencing and contrasts tested in the differential expression analyses. (**A**) Large single cells were isolated to form monoclonal vegetative cultures. These were manipulated in size to drop below the sexual size threshold (SST) and significantly reduce life cycle duration. In such a manner cells above and below SST of the same genotype were cultured simultaneously and harvested for RNA-seq. Cells are coloured to represent the different mating types and size classes, with the line between two shades representing the raphe. (**B**) Cell lengths at the time of harvesting for RNA-seq experiment. (**C**) Transcriptomic differences between mating types, size classes and their interaction were studied by differential expression analysis over four different contrasts. (**D**) Bar plots showing the number of differentially expressed *C. closterium* genes for each contrast, each time subdivided in genes upregulated in group A vs group B, and genes upregulated in group B vs group A.

Protein domain and gene family annotation of *C. closterium* genes was performed using TRAPID 2.0 (Bucchini *et al*., 2021), with default parameters (E-value threshold 10^-5^) and PLAZA Diatoms as a database for similarity searches. Meanwhile, InterPro and gene family functional annotation for *S. robusta* was retrieved from PLAZA Diatoms 1.0 (Osuna-Cruz *et al*., 2020). Additionally, computational predictions of subcellular localization, phylostratigraphic gene age as well as enrichment analyses were performed, as detailed in the **Supplementary Methods**.

### Reverse transcriptase quantitative PCR (RT-qPCR) analysis of selected candidate genes

An RT-qPCR experiment was performed on six genes (Ccl_4437, Ccl_5484, Ccl_13556, Ccl_14859, Ccl_14861 and Ccl_16260) to validate results of the size-vs-mating type RNA-seq experiment. An identical set-up as for the RNA-seq experiment was used to prepare the cDNA of four strains (A5, C2, MC3, MC5), two biological replicates for each mating type, in two size variants above and below the SST.

Details about cDNA preparation, primers (**Supplementary Table 2**), cycling conditions and statistical analysis of the results can be found in **Supplementary Methods**.

### Transcriptomic response to sex-inducing pheromone minus (SIP-)

Further, we performed an RNA-seq experiment to specifically asses the response of three MT+ specific genes (Ccl_14859, Ccl_14861 and Ccl_14874 to sex-inducing pheromone of the opposing mating type. Prior to the experiment SIP containing medium from MT-strain CA1.15 (MT-, DCG 0923) was harvested as described previously (Klapper *et al*., 2021). The activity of the filtered medium was tested by its ability to induce a partial cell cycle arrest after 6 h of treatment in two biological replicates of MT+ strain ZW2.20 using an Amnis ImageStreamX Mk II imaging Flow Cytometer (Luminex), following the protocol of Audoor *et al*., 2024 (**Supplementary Figure 3**). Next, cultures of MT+ strain ZW2.20 (MT+, DCG 0922) growing in the mid-exponential phase were placed in the dark for 24 h. Two hours before re-illumination, the supernatant medium of ZW2.20 was replaced with either filtered medium from its mating partner CA1.15 (treatment) or fresh culture medium (control). Triplicate ZW2.20 cultures were harvested at two time points (3 h, 9 h) after re-illumination for both control and SIP treatment conditions. Culture harvesting and RNA extraction was performed as explained above. Library preparation (including poly-A enrichment) and paired-end Illumina 150 bp mRNA sequencing was performed at Novogene (UK) Co., Ltd. Details about quality trimming, mapping and differential expression analysis of the results can be found in **Supplementary Methods**.

### Phylogenetic analysis of MRM2 homologs

A hidden Markov model (HMM) was constructed to define the MRM2 gene family (HOM02SEM009752) and identify more distant homologs. A protein sequence alignment created with MAFFT v7.453 (Katoh & Standley, 2013) was trimmed to retain only the conserved LRR domain, after which a profile HMM was created using the hmmbuild command of HMMER (hmmer.org). The profile HMM is included in **Supplementary Data** as a reference for the MRM2 family. Next, a targeted search for MRM2 homologs in eukaryotic genomes was performed with the hmmsearch command and the top-50 proteins with the highest domain-specific E-value score were selected for phylogenetic analysis. A detailed pipeline can be found in **Supplementary Methods**.

### Data availability

New RNA-seq libraries generated for *C. closterium* and *S. robusta* are available at ENA with project number PRJEB52586 (size and mating type) and PRJEB81513 (SIP response). Differential expression calls for both *C. closterium* and *S. robusta* as well as gene family predictions for *C. closterium*, curated gene models of MRM2 family proteins in selected pennate diatoms, and the MRM2 family HMM model are available as **Supplementary Data**.

## Results & Discussion

### Transcriptome profiling reveals size- and mating type-dependent gene expression

We profiled gene expression of large- and small-sized cells for three independent *C. closterium* cultures of each mating type. To identify only genes that show a robust response across strains, three different genotypes of MT+ (C2, MC4, MC5) and MT-(A5, A6, MC3) each were used as biological replicates. Only genotypes that belong to *C. closterium* clade II were selected, to avoid sequence divergence affecting the mapping to the reference genome (CA1.15, clade II) (Stock *et al*., 2019; Audoor *et al*., 2024).Small cells were generated by microsurgery of the apices of large cells, shortening the time required for cells to become sexually inducible (**Supplementary Figure 2**). Indeed, we estimate that more than one year of optimal growth (with a daily size decrease of ∼0.11 µm) would be necessary for cells with an initial cell size of around 100 µm to naturally drop below the SST, which we determined to occur at a cell size between 55 - 60 µm, unlike the previously reported 66µm (Vanormelingen *et al*., 2013). (**Figure 2A**). Using the microsurgery technique, we obtained matched cultures for each genotype in two size variants, before and after size manipulation. This eventually resulted in paired cultures with sizes on average 76±7µm (>SST) and 36±8µm (<SST) that were harvested for RNA-seq (**Figure 2B**). All cultures were tested for their ability to engage in sexual reproduction prior to the experiment. In none of the large-celled cultures, any stage of sexual reproduction (paired cells, auxospores, initial cells) could be identified after mixing compatible mating types, while all sexual stages were observed in all cultures whose cell size was experimentally decreased.

To desynchronize the cell cycle and thus prevent the enrichment of a particular cycle phase cultures were maintained in constant light regime for at least four weeks. RNA-seq reads of all samples were mapped to the reference genome of *C. closterium* strain CA1.15 (MT-) (Audoor *et al*., 2024). The average mapping rate was 62.5 %, with no significant differences in mapping rate between large (63.1±8.0%) and small cells (61.9±4.9%) or between MT+ (62.2±7.0%) and MT-(62.9±6.4%) (p>0.05, two-sample t-test). Multidimensional scaling showed clustering of samples by parental genotype (A5, A6, C2 offspring of CA1.15xZW2.20 versus MC3, MC4, MC5, offspring of NH3.13xWS3.7) and not by condition (size or mating type), in agreement with recent observations of significant genotype-specific differences in expression patterns of the centric diatom *Skeletonema marinoi* (Pinseel *et al*., 2022) (**Supplementary Figure 4**).

Differential expression (DE) analysis was performed over four contrasts to detect genes responsive to cell size, mating type, or the interaction of cell size and mating type (**Figure 2C**). Overall, a set of just 112 out of 24.633 annotated genes was identified as differentially expressed in one or more of these contrasts (**Supplementary Data**). Given that these DE genes show a conserved expression pattern throughout three independent genotypes for each mating type, we expect a significant role for these genes in the size- and mating type-related response in *C. closterium*. The small number of DE genes suggests that transcriptomic differentiation between cell size classes and mating types in diatoms is limited to specific traits related to size sensing and behavioral sexual dimorphism, while the overall metabolism of cells remained unchanged. RT-qPCR was performed on independent cultures belonging to both mating types and size classes for validation, confirming the mating type- and size-dependent expression of five genes (Ccl_14859, Ccl_14861, Ccl_16260, Ccl_4437 and Ccl_13556) while failing to find a response for Ccl_5484 (**Supplementary Figure 5**).

Whereas 21 genes were upregulated in small cells regardless of their MT, no genes were upregulated in large cells. The cellular changes that take place in both mating types when cells pass the SST thus appears to be mainly driven by the activation of genes below rather than by suppression of genes expressed above the SST. LWe extended the search for size-related genes by performing a paired analysis in which the effects of genotype were subtracted for each corresponding large-vs-small pair (**Supplementary Data**). In this way, we identified a larger set of 182 differential expressed genes, comprising 169 genes upregulated in small cells, and 13 genes that were biased towards large cells.

A number of genes was differentially expressed between mating types regardless of their size: 8 and 54 genes were induced in MT- and MT+, respectively (**Figure 2D**). Hence, the two mating types can be distinguished by their unique gene expression profiles throughout their entire life cycle, i.e. across large and small stages. Once cells decrease below the SST, an additional activation of mating type specific genes occurs. Indeed, 29 genes were upregulated in cells of a specific mating type but only below the SST, with a marked predominance of MT+ specific genes (26 versus only 3 MT-specific) (**Figure 2D**). Hence, size-dependent gene expression changes also reflect the physiological and behavioral differentiation between mating types, even in the absence of a compatible mating partner.

### Regulatory genes and retrotransposon activity mark small and MT+ cells respectively

For 72 of the differentially expressed genes, InterPro protein domains and/or PLAZA Diatoms gene families could be assigned, whereas 40 genes in our dataset remained without functional annotation (**Supplementary Data**). Enrichment tests were performed to identify InterPro domains and gene families that were significantly more common among the DE genes compared to the background of all expressed genes. We detected eight gene families and five InterPro domains that were significantly enriched among the DE genes for specific contrasts (**Figure 3**). Moreover, one additional gene family was enriched in the set of all DE genes.

**Figure 3:**
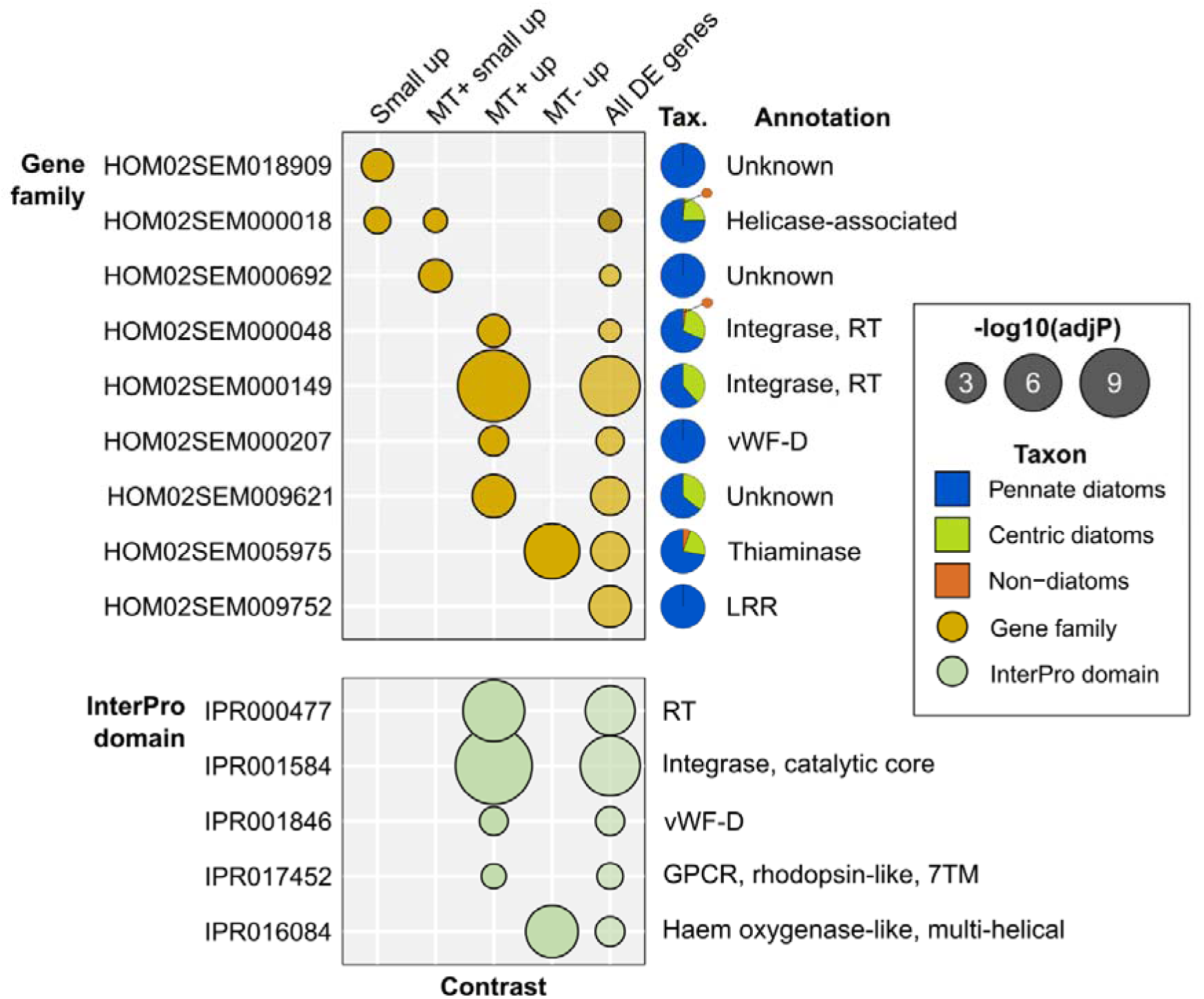
Enrichment of gene families and protein domains among differentially expressed (DE) genes from different contrasts (x-axis). Features that are significantly enriched in a contrast are shown by circles, whose size scales with the significance level (-log10). Pie charts show the taxonomic composition of gene families. MT: mating type, RT: reverse transcriptase, PDE: phosphodiesterase, GPCR: G-protein coupled receptor, LRR: Leucine-rich repeat, vWF-D: Von Willebrand factor type D, TM: transmembrane domain.

Genes upregulated in small cells of both mating types appear to be dominated by regulatory functions. These include a family encoding DNA-binding helicase-like proteins (HOM02SEM000018) (**Figure 3**). Several other genes that interact with DNA or RNA were upregulated in small cells: a nuclease (Ccl_23878), a 5’-Nucleotidase (Ccl_12722) and a ribonuclease H (Ccl_21492). Furthermore, five genes upregulated in cells below the SST contain a potential nuclear localization signal, among which two genes from the family of unknown function HOM02SEM018909. The identification of putative regulatory genes that are transcribed in both mating types below the SST is especially relevant in light of recent reports that the sex-determining gene *MRP3* of *P. multistriata* and a methyltransferase situated nearby are differentially methylated between mating types (Ruggiero *et al*., 2023). A general size-responsive transcription factor is thought to preferentially activate the non-methylated sex-determining allele, causing mating type differentiation below the SST (Ruggiero *et al*., 2023). Hence, the nuclear and DNA-binding genes identified here are prime candidates as size-dependent molecular switches in diatoms.

Two families of proteins containing reverse transcriptase and DNA integrase domains, indicating a retrotransposon origin, were significantly enriched among MT+ biased genes: HOM02SEM000149 and HOM02SEM000048 (**Figure 3**). Indeed, reverse transcriptase and integrase InterPro domains were significantly more abundant among MT+ biased genes than in the rest of the expressed transcriptome (**Figure 3**). The role of retrotransposons in the evolution of mating type identity may be similar to mechanisms observed in other eukaryotic groups where the mating type locus is protected through suppression of recombination caused by retrotransposon accumulation (Bachtrog, 2006; Kent *et al*., 2017). Alternatively, an overall higher activity of (former) transposable elements in MT+ may be the result of differences in chromatin accessibility between mating types similar to the above-mentioned differential methylation in *P. multistriata* and as is the case in sex chromosomes of higher animals and plants (Slotkin & Martienssen, 2007; Hobza *et al*., 2017), or may be the result of random differences in transposon activity between selected genotypes for each mating type.

### A gene cluster of MT+ and small size specific genes plays a role during sex pheromone signaling

Among the differentially expressed genes for size and mating type, we identified three *C. closterium* genes that are located nearby on contig81: Ccl_14859, Ccl_14861 and Ccl_14874 (**Figure 4A**). Gene expression of each of the three genes was extremely specific for small cells and restricted to the MT+ (**Figure 4A**). Consulting publicly available RNA-seq datasets, we found that the three aforementioned genes were on average 109-fold upregulated during early sexual reproduction in available RNA-seq data on top of their baseline expression in small cells (**Figure 4A**) (Audoor *et al*., 2024). Inspecting the expression pattern following the crossing of compatible strains ZW2.20 and CA1.15 (T1: 9 h, T2: 14 h, T3: 27 h), we observed that the induction of Ccl_14859 and Ccl_14861 was the strongest at the first time point T1, which is the stage when sex pheromone signaling, mate finding and mating pair formation is most intense (**Figure 4A**). The expression of Ccl_14874 peaked somewhat later: between stage T1 and T2, when gametogenesis is at its maximum. Since the rapid induction of these three genes during the early stages of mating may indicate a role in the response to sex inducing pheromone (SIP) of the opposite mating type (Klapper *et al*., 2021), we set up a time-series RNA-seq experiment (**Figure 4B**). We specifically focused on the interval during which cells become conditioned by SIP before the formation of mating pairs, by treating MT+ cells with SIP-containing filtrate and assessing global gene expression after 3 h and 9 h of re-illumination following a dark-arrest. Compared to the untreated control, the expression of each of the three genes was significantly induced by a treatment of 9 h with SIP-containing filtrate (**Figure 4B**). Compared to their initial expression in small MT+ cells, both Ccl_14861 and Ccl_14874 were stimulated about 30-fold after 9h.

**Figure 4:**
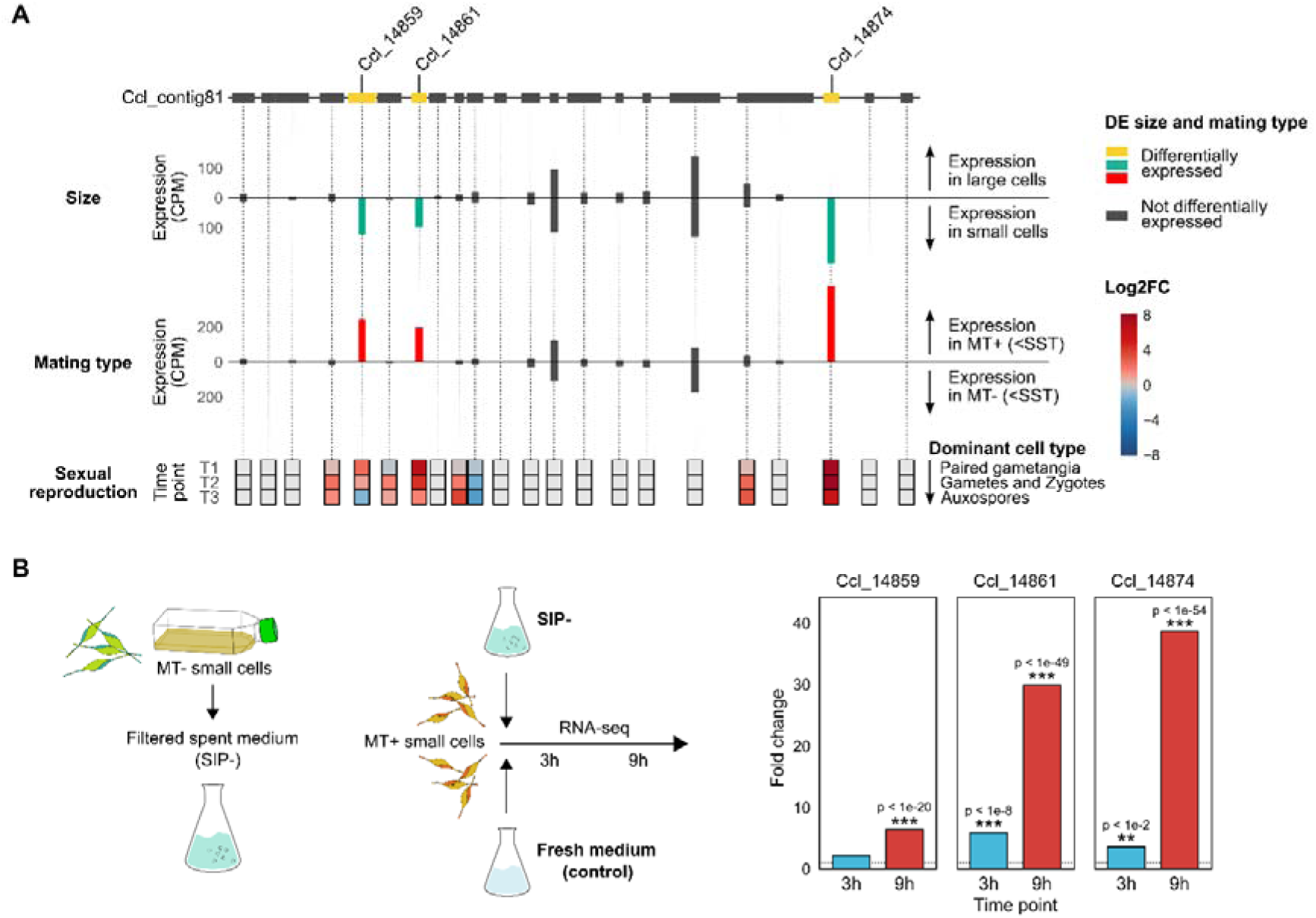
Gene expression of adjacent genes on *C. closterium* contig81 in different size classes, mating types and during three stages of sexual reproduction. (**A**) Bar plots show the average expression over replicates. The expression of individual replicates is visualized in Supplementary Figure 8 and Supplementary Table 3. Heatmaps show the log2 fold change between sexual crosses and vegetative control cultures across three time points: T1 (9 h post-cross, dominated by paired gametangia), T2 (14 h post-cross, gametes and zygotes) and T3 (27 h post-cross, auxospores). Expression data of sexual reproduction was obtained from Audoor *et al*., (2024). CPM: counts per million, MT: mating type, SST: sexual size threshold, log2FC: log2(fold changes). (**B**) Bar plots show fold changes after 3 h and 9 h of re-illumination following a treatment of MT+ cells with Sex Inducing Pheromone (SIP) containing spent medium. Fold changes represent the expression after SIP treatment relative to the matched untreated control. Stars indicate the significance of differential expression between SIP treated and control conditions for each gene in each time point (false discovery rate-adjusted p-values, * p<0.05, ** p<0.01, *** p<0.001).

In terms of functional annotation, the amino acid sequence of Ccl_14859 shares similarity with the *P. tricornutum* phytotransferrin Phatr3_J54465 (BLASTp against PLAZA Diatoms v1.0 database, E-value: 3e-88, **Supplementary Figure 6**) (McQuaid *et al*., 2018; Turnšek *et al*., 2021) and encodes an N-terminal signal peptide. The other two genes (Ccl_14861, Ccl_14874) which differ only by two amino acids at positions 303 and 306 (**Supplementary Figure 7**), had no functional annotation except for an N-terminal signal peptide, indicating a secretory pathway. Furthermore, no homologs could be detected in any other diatom genome in the PLAZA Diatoms v1.0 database (BLASTp, E < 1e-3), making these two genes highly taxon-specific markers for MT+ identity.

The presence of three closely located genes with a shared MT+ specific and SIP-responsive properties suggests a shared function during early mate finding. While functional gene clusters are commonly found in other eukaryotic taxa such as plants, animals and fungi (Nützmann *et al*., 2018), where they play a role in the same cellular process, often a metabolic pathway, the only clear example known in diatoms has been described for fucoxanthin chlorophyll proteins (Devaki & Grossman, 1993). Co-expression of clustered genes is typically a result of co-regulation by the same transcription factors, through shared binding sites or common chromatin-level regulation (Nützmann *et al*., 2018). Further studies are warranted to elucidate the functional significance and regulatory elements associated with this gene cluster.

### Mating type differentiation and size sensing genes are associated with pennate- and *Cylindrotheca*-aged genes

Phylostratigraphic analysis, a method used to date the origin of genes or gene families by looking at their homologs across different species, revealed that size- and mating type-responsive DE genes were on average evolutionarily young compared to the entire expressed transcriptome. Indeed, genes upregulated in small cells, in one or both mating types, were significantly enriched in genes for which we could not detect homologs outside the *C. closterium* genome (**Figure 5A**), and the same was true in the size-dependent genes from the additional paired analysis (adjusted p-value < 1e^-5^, hypergeometric test). These genes are most likely involved in the species-specific pheromone signaling and mate finding behavior, which become only apparent below the SST (Klapper *et al*., 2021).

**Figure 5:**
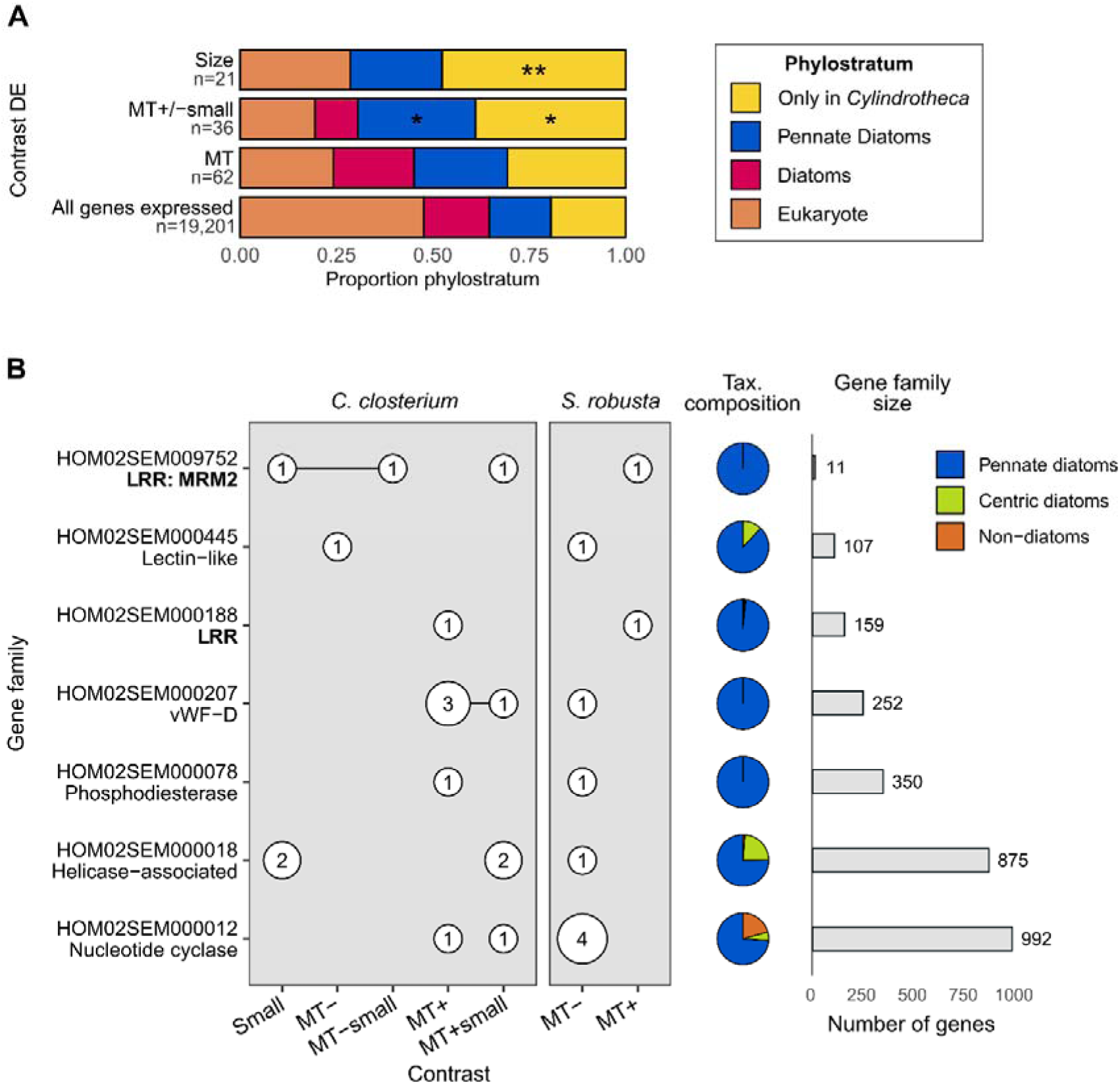
Phylostratigraphic analysis of size and mating type-responsive genes and comparative analysis of gene families containing genes with a mating type- or size-biased expression in two pennate diatoms: *Cylindrotheca closterium* and *Seminavis robusta*. (**A**) Bar plots showing the proportion of different phylostrata (phylogenetic gene ages) among sets of differentially expressed (DE) genes as well as the background (all expressed genes). Stars indicate the enrichment significance level of a given phylostratum relative to the background (*: p < 0.05, **: p < 0.01, hypergeometric test), MT: mating type. (**B**) For each gene family, the number of differentially expressed genes in each contrast is shown by circles, with genes that were differentially expressed in more than one contrast connected by a horizontal line. Pie charts show the taxonomic composition of gene families, while bar charts show the size (number of genes) of each family. LRR: leucine-rich repeat, vWF-D: Von Willebrand factor type D domain.

Size-sensing and size-dependent sexual maturation have been observed for most diatom species from both centric and pennate clades (Chepurnov *et al*., 2004; Poulíčková *et al*., 2019). Nevertheless, none of the 21 purely size-responsive genes we found in *C. closterium* were of diatom age, i.e., whose gene family was restricted to pennate and centric diatoms (**Figure 5A**). While such diatom-aged size genes may have been missed due to the large variability between genotypes limiting the power to detect DE genes, it may also suggest that size-sensing in diatoms does not share a common master regulator. In other eukaryotic organisms, physicochemical signals such as the scaling of the nucleus and membrane-free structures, i.e., nucleolus, spindle, and microtubules, have been used to measure their size in comparison to the cell size, the cell volume or the surface area (Marshall, 2015; Reber & Goehring, 2015). Hence, mating type differentiation and sexual maturation may proceed directly downstream of such physical signals through intracellular scaling without a transcriptionally regulated master regulator.

Genes that were differentially expressed between mating types but only in small cells were significantly more likely to be of pennate age compared to all other *C. closterium* genes (**Figure 5A**). Indeed, many (30%) of the genes involved in mating type differentiation below the SST were rooted within the pennate diatoms. Given that heterothally is a common reproductive strategy in pennate diatoms while centric diatoms typically display homothallic reproduction without mating types (Poulíčková *et al*., 2019), these results suggest that the molecular mechanisms underlying mating type determination have a shared origin in the pennate diatoms around 137 million years ago (Nakov *et al*., 2018).

### Leucine-rich repeat proteins are involved in mating type differentiation in pennate diatoms

We next tried to pinpoint which common genes drive mating type differentiation in pennate diatoms by performing a comparative analysis using existing transcriptomic libraries from another heterothallic pennate diatom, *Seminavis robusta* (Osuna-Cruz *et al*., 2020). We took advantage of the presence of 36 and 12, vegetative RNA-seq samples from small untreated MT+ and MT-cells respectively, to perform a differential expression analysis. This resulted in a set of 183 genes that showed a mating type-specific expression response. We intersected the gene families of these *S. robusta* mating type-specific genes with families to which the 112 *C. closterium* size- and mating type-biased genes belong. Seven gene families contained DE genes in both species (**Figure 5B**).

Although the seven shared gene families range from small (11 genes) to large (992 genes), all of them are predominantly composed of genes belonging to pennate diatoms, further corroborating how mating type-dependent genes have evolved simultaneously with heterothally at the origin of pennate diatoms. Genes from two families containing Leucine-rich repeat (LRR) InterPro domain proteins showed a mating type biased response in both *C. closterium* and *S. robusta* (**Figure 5B**). Since the LRR domain is typically involved in protein-protein interaction or signaling, their conserved mating type dependent response suggests a role in communication between the opposite mating types in pennate diatoms. Although the first LRR family HOM02SEM000188 includes one *S. robusta* and one *C. closterium* gene with a MT+ specific expression, phylogenetic analysis showed that these two genes are not orthologous (**Supplementary Figure 9**). Hence, the functional and evolutionary significance of their shared expression pattern is unclear.

On the other hand, HOM02SEM009752, a pennate-specific family with only 11 genes in 8 species, comprises two *C. closterium* genes, both of which are specific for small cells, but with a contrasting expression pattern between mating types: gene Ccl_4437 was upregulated in MT+ cells, whereas Ccl_5484 was upregulated in MT-cells (**Figure 5B**). The *S. robusta* homolog (Sro702_g189950) showed a strong MT+ specific expression. Importantly, this family encodes the Mating-type Related Minus 2 (*MRM2*) gene that was previously identified in the raphid pennate diatom *P. multistriata*. *MRM2* is a component of the mating type determination system of *P. multistriata*, where it is downstream of the sex-determining gene *MRP3* (Russo *et al*., 2018).

Importantly, a paired analysis identified three LRR genes (Ccl_1352, Ccl_11832, Ccl_7129) that are upregulated in small cells irrespective of the mating type, suggesting that some LRR proteins may be involved in bidirectional recognition between mating types below the SST.

### Mating-type Related Minus 2 (MRM2) is widely conserved in pennate diatoms

Multiple sequence alignment showed that proteins from the MRM2 family are conserved through their LRR domain. Therefore, we set out to identify additional homologs of MRM2 by constructing a profile hidden Markov model (HMM) of the LRR domain in the 11 proteins affiliated with HOM02SEM009752. Subsequently, the profile HMM was used to search the PLAZA Diatoms reference dataset for additional proteins that are homologous to MRM2 through their LRR domain (Osuna-Cruz *et al*., 2020). Phylogenetic analysis based on the full-length protein sequence of the top-50 hits revealed a well-resolved clade containing the original MRM2 gene family as well as two genes belonging to other gene families (**Figure 6A, Supplementary Figure 10**). The proteins in this MRM2 clade share structural similarities, consisting of a predicted transmembrane domain on the C terminus and the LRR domain displayed extracellularly (**Figure 6A**). Motif discovery in the LRR domains of MRM2 homologs revealed the NxLt/sGxIPx motif typical to the plant specific (PS) family of diatom LRR proteins (**Figure 6A**) (Schulze *et al*., 2015). The short (∼50AA) N-terminal intracellular domain does not contain a kinase domain or other known signaling functionality, similar to the plant LRR-RLPs (leucine-rich repeat receptor-like proteins), which associate with co-receptors to transduce immune signals to the plant cell (Snoeck *et al*., 2023). Together, MRM2 homologs were found in the genome of seven raphid pennate diatoms, as well as the araphid *Synedra acus*, suggesting a common origin at the root of the pennate diatoms.

**Figure 6:**
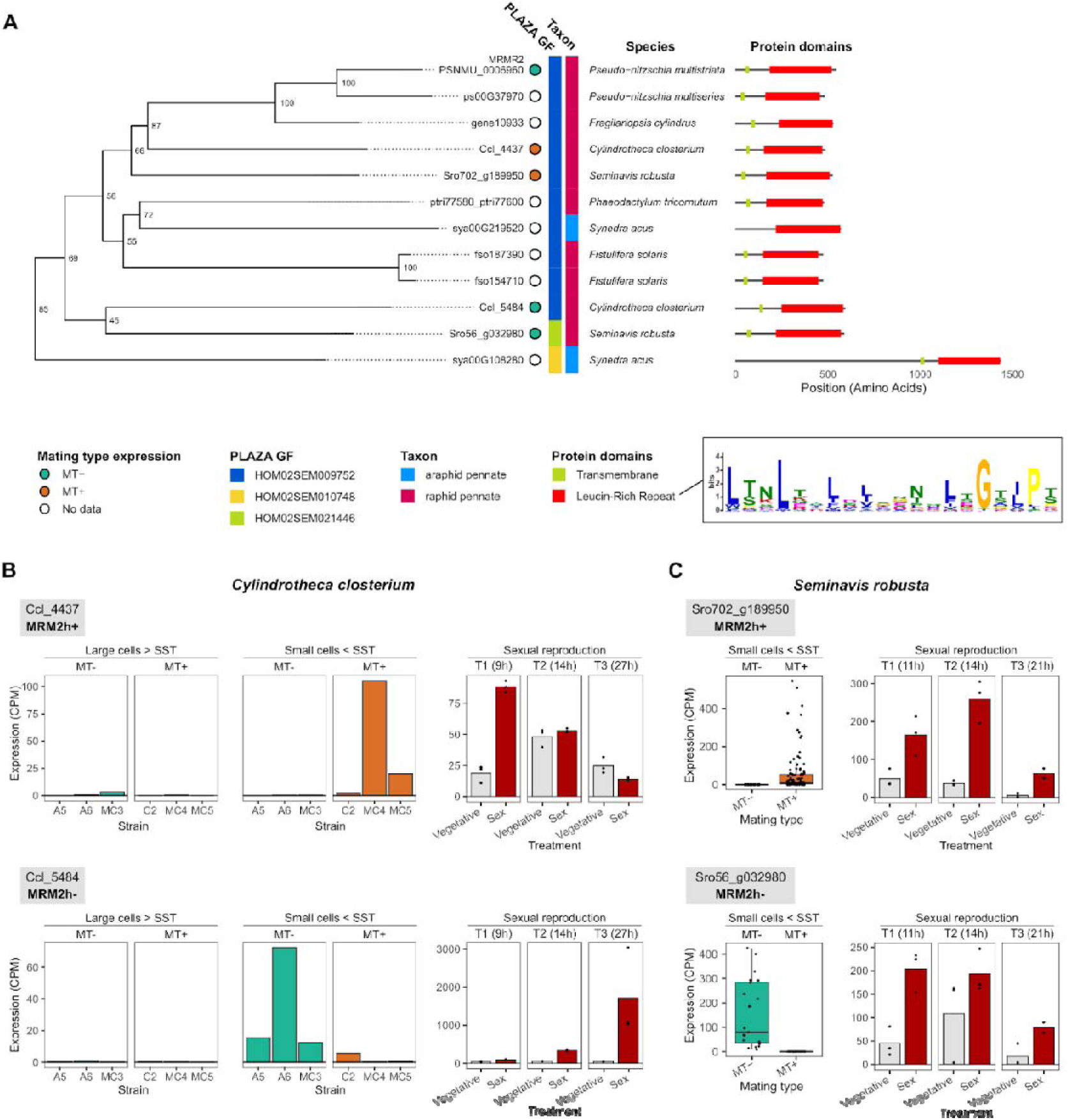
Phylogeny and expression of MRM2 homologs in pennate diatoms. **(A)** Bootstrap-consensus phylogenetic tree of MRM2 proteins in diatoms. Bootstrap support is shown as node labels. The full phylogenetic tree of the top-50 best MRM2 hits is available as Supplementary Figure 10. Next to the gene labels, their mating type-dependent expression pattern, gene family (GF), taxonomy and protein structures (InterPro domains and transmembrane domains predicted by Phobius) are shown. The inset shows the MEME sequence logo of the leucine-rich repeat domains of these proteins. **(B)** Bar plots showing expression in counts per million (CPM) of MRM2 homologs in large and small cells of different *Cylindrotheca closterium* strains (left) and during three stages of sexual reproduction (right). For the sexual reproduction, bar plots show the average expression while dots show the individual RNA-seq samples. SST: sexual size threshold, MRM2h+: MT+ biased homolog of MRM2, MRM2h-: MT-biased homolog of MRM2. **(C)** Expression of MRM2 homologs in the *S. robusta* expression atlas (Osuna-Cruz *et al*., 2020). Left: boxplots of expression in MT+ (n = 108) and MT-(n = 21) samples in a range of non-sexual conditions, spanning biotic interactions, environmental conditions, sexual reproduction, and diel cycles. Right: expression during sexual reproduction covering three time points after crossing. Bar plots show average expression while dots show individual RNA-seq samples.

### Duplicated homologs of *MRM2* show an inverted mating type identity

The *MRM2* homologs of *C. closterium* and *S. robusta* have seemingly undergone gene duplication, with Ccl_4437 and Sro702_g189950 being closest related to the *P. multistriata MRM2* gene, whereas Ccl_5484 and Sro56_g032980 are more distant (**Figure 6A**). These distant out-paralogs of *MRM2* are lost or missing in most other pennate diatoms that encode only one copy of *MRM2*, including the original gene identified in *P. multistriata* (Russo *et al*., 2018).

Our data suggest that both Ccl_4437 and Ccl_5484 are conditionally expressed only in cells below the SST (**Figure 6B**). Ccl_4437 was specifically expressed in MT+ strains, while Ccl_5484 showed the opposite expression pattern, being biased towards MT-. We propose calling these homologs *MRM2h*+ (*MRM2* homolog expressed in MT+) and *MRM2h*-respectively. Expression profiles of the small cells from the *S. robusta* expression atlas are congruent with what we observed in *C. closterium*: one homolog (Sro702_g189950, *MRM2h*+) is MT+ specific, while the other (Sro56_g032980, *MRM2h*-) is MT-specific (**Figure 6C**). To explore the variability of this response in other genotypes beyond the ones included in the *S. robusta* expression atlas (MT-: 84A, 85B, MT+: 85A), we performed RNA sequencing of six additional *S. robusta* genotypes below the SST, three for each mating type (MT+: 85A-9, P5, P7 and MT-: 85A-1, 85A-6, P4). While this experiment confirmed the MT-specific expression of *MRM2h*-in two out of three strains, *MRM2h*+ was expressed at a very low level in all six strains (average expression of 0.76 counts per million). Hence, the mating type specific expression pattern of *MRM2h*+ could not be confirmed in these additional strains (**Supplementary Figure 11**).

Combined, the mating type specific expression of *MRM2* homologs in three species (*P. multistriata, C. closterium* and *S. robusta*) and its distribution throughout the pennates point to a role for *MRM2* as a key gene for mating type identity. As transcription of *MRM2* homologs was activated only when the cell size decreased below the SST, the function of MRM2 appears to be specific for sexually competent cells. Similar studies on the transcriptomic response to cells passing the SST should be undertaken in other life cycle model organisms such as *S. robusta* and *P. multistriata* to determine if MRM2’s size-dependency is conserved throughout the pennate diatoms. The duplication and inversion of mating type preference in *S. robusta* and *C. closterium* suggests that MRM2 homologs have evolved to play similar roles, yet in opposite mating types. Finally, our data encourages discussion about how mating type identity is assigned. Mating types are currently distinguished based on cellular behavior during mating: MT-represents passive cells, while either the MT+ gametangia or gametes are motile (Kaczmarska *et al*., 2013). Our data suggests that distinguishing mating types by their behavior may not reflect molecular evolution. To illustrate, *MRM2h*+ genes Ccl_4437 and Sro702_g189950 exhibited MT+ expression but are phylogenetically closest to the original *P. multistriata MRM2*, which was expressed in MT-(**Figure 6**).

### Upregulation of *MRM2* homologs during sex suggests a function in recognition between compatible mating types

The differentiation between mating types in raphid pennate diatoms is most apparent during sexual reproduction, where attraction and cell-cell recognition ensure the correct pairing of compatible gametangia and fusion of a MT+ with a MT-gamete (reviewed in Bilcke et al., 2022). Therefore, we inspected the expression of *MRM2* homologs during different stages of sexual reproduction (Audoor *et al*., 2024; Osuna-Cruz *et al*., 2020) (**Figure 6B,C**). Both *C. closterium MRM2* homologs as well as the *S. robusta MRM2h*+ were significantly transcriptionally upregulated in at least one stage of sexual reproduction. Strikingly, the *C. closterium* gene *MRM2h*+ is only upregulated during the early stage whereas *MRM2h*-was most strongly upregulated in the last stage. Hence, not only are *MRM2* homologs in *C. closterium* specific for the opposing mating types but they also appear to play a role in different stages of sexual reproduction: gametangia pairing (*MRM2h*+) versus the differentiation of gametes or auxospore development (*MRM2h*-).

## Conclusions and Outlook

In this study, we leveraged experimental cell size manipulation to decipher the transcriptional reprogramming associated with size-based sexual maturation in diatoms. By simultaneously harvesting the same six genotypes in large and small cells, we avoided potential confounding due to the use of different genotypes in the two size classes. Furthermore, the use of full biological replication (i.e. independent genotypes) further ensures that the small set of DE genes we identified are robust and valid for a range of *C. closterium* strains. In general, size and mating type were not associated with large transcriptional differences in our experiment, which resulted in relatively few DE genes (< 100 genes per contrast). This may suggest that only few genes are involved in the differentiation of size and mating type classes or may alternatively be caused by the large variability between genotypes, making it harder to detect true DE genes. Indeed, correcting for the genotype effect in a paired analysis increased the number of identified genes by about 10-fold (from 21 to 182 genes), but this was only possible for the “Size” contrast.

A set of 29 mating type-specific genes were transcriptionally activated in cells that have passed the SST. We argue that such genes must underlie the chemical and behavioral differentiation between mating types, which only becomes apparent in small cells. Three such co-expressed genes were located nearby on the same contig. Their confined expression in small MT+ cells and coordinated induction in response to sex pheromones suggest an important role during the pairing process, in which MT+ cells are actively gliding towards an attracting MT-cell.

Furthermore, we demonstrated that the *P. multistriata* gene *MRM2* not only has a wider distribution in pennate diatoms than was previously reported (Russo *et al*., 2018), but also displays mating type-specificity in at least three raphid pennate diatoms from diverse taxonomic clades (*C. closterium, P. multistriata, S. robusta*). Judging from the fact that these plant-like LRR receptor proteins are most strongly expressed during cell pairing and gamete fusion, we argue that MRM2 is part of an ancient partner recognition system that dates back to the origin of the pennate diatoms. Consequently, the conservation of MRM2 is the first molecular evidence for heterothally being an ancestral feature in pennate diatoms. While most pennate diatoms indeed display a heterothallic mating system, some pennate species display homothallic reproduction or facultative heterothally (Poulíčková *et al*., 2019). Heterothally would consequently have been secondarily lost in these species. Additionally, in accordance with Russo and colleagues (2018) we did not find any homologs of the *P. multistriata* mating locus *MRP3* outside the *Pseudo-nitzschia/Fragilariopsis* clade (Russo *et al*., 2018), suggesting that different clades use different mating type determination systems. In future studies, assessing the conservation of mating type dependent genes such as *MRM2* in a diverse set of heterothallic and homothallic pennate diatoms will be instrumental to understand their evolutionary history. In that respect, the 100 Diatom Genomes project (<u>jgi.doe.gov/csp-2021-100-diatom-genomes</u>) will soon make it possible to investigate life cycle genes at an unprecedented scale.

While the earliest reports of the unique size reduction-restitution life cycle of diatoms date as far back as 1869 (Pfitzer, 1869; Macdonald, 1869), the mechanism that cells use to accurately measure their own size is still completely unknown. The fact that the size is constantly monitored by the cell can be deduced from the sexual maturation after the experimental cell size manipulation we performed using a glass Pasteur pipette. Such microsurgery-based size manipulation, which was previously executed using a razor blade by van Stosch in 1965 (von Stosch, 1965) and later repeated by Roshchin (Roshchin, 1994) and Chepurnov (Chepurnov *et al*., 2004), represents a time and cost-efficient way to significantly reduce the generation time of experimental models such as *C. closterium*. Using this approach, we identified a set of 21 genes that are upregulated in small cells (<SST) relative to large cells (>SST) in both mating types. These size markers include several nucleus-localized and DNA-interacting genes, suggesting a function as size-dependent switches, activating downstream sexual maturation genes only below the SST, but need to be further validated by functional experiments. However, none of these genes could be clearly linked to a physical function in cell size sensing, and neither did any of them occur specifically in the genome of both pennate and centric diatoms. Hence, it remains unexplained how cells measure their own size and whether a diatom-wide transcriptional master regulator exists, which our analysis may have lacked the power to detect. The identification of the first molecular markers for cell size dependent sexual maturation in diatoms - including conserved key genes such as *MRM2* - paves the road for future research into the physicochemical and molecular changes taking place when cells pass the SST.

## Acknowledgements

G.B. is a postdoctoral fellow supported by Fonds Wetenschappelijk Onderzoek (FWO, 1228423N). W.V. acknowledges partial financial support from FWO project G001521N, BOF/GOA projects No. 01G01715 and 01G01323, as well as infrastructure funded by EMBRC Belgium-FWO project GOH3817N. The BCCM/DCG culture collection is supported by the Belgian Science Policy (Belspo).

We sincerely thank Dr. Koen Van den Berge for helpful discussion about experimental design. We are grateful towards Olga Chepurnova for help with maintaining and manipulation of diatom cultures in cryopreservation tanks. We would further like to thank Dr. Katerina Pargana for her help with bioinformatic analysis and Dr. Petra Bulánková for assisting with molecular biology protocols.

## Competing Interests

The authors declare that they have no conflict of interest.

## Authors Contribution

DB, GB, and WV conceived the study with input from KV and LDV. DB performed microsurgery and transcriptomic experiments. DB and SD jointly performed RNA extraction and quality control, while SA and DB were responsible for RT-qPCR of selected genes. Differential expression and comparative transcriptomic analyses were performed by GB with the support of DB. DB, GB, WV, and KV wrote the manuscript. All authors commented on and approved of the manuscript.

## Notes

### Competing Interest Statement

The authors have declared no competing interest.

### Summary of Updates

The manuscript has been revised, and several changes have been implemented. New data and supporting information have been added to the results and the supplementary data. Detailed information about microsurgical removal of cell apices has been included. RT-qPCR showing overexpression of SIP (sex-inducing pheromone) responsive genes has been substituted by RNA-seq data. Paired differential expression analysis of the RNA-seq data has been added.

