## Supplementary Methods for "Molecular fingerprints of cell size sensing and mating type differentiation in pennate diatoms"

Generation of *Cylindrotheca closterium* and *Seminavis robusta* genotypes, and size manipulation of *C. closterium* cells

The original parental strains of *C. closterium* CA1.15 (MT-) and NH3.13 (MT-) were crossed in sterile Petri dishes (PS, 60/15 MM, with vents, Greiner BIO-ONE) with ZW2.20 (MT+) and WS3.7 (MT+) respectively (**Supplementary Table 1**). In the case of *S. robusta*, the parental strain Ponton 36 (MT+) was mated with VM3-32 (MT-), and 85A (MT+) with VM3-12 (MT-), resulting in progeny strains shown in **Supplementary Table 1**. For both species, single progeny cells were isolated using an Olympus® CKX41 Inverted Microscope and a glass Pasteur pipette and placed into a 96-well flat bottom plate (Greiner CELLSTAR®, Sigma Aldrich Belgium) where they propagated in monoclonal progeny strains. Two weeks after isolation, cultures were transferred to a 24-well plate (Greiner CELLSTAR®, Sigma Aldrich Belgium), where they were maintained prior to further experiments. A subpopulation of each *C. closterium* strain was transferred to sterile a Petri dish (PS, 60/15 MM, with vents, Greiner BIO-ONE), where cells were submitted to manual manipulation of cell size using a glass Pasteur pipette with an ultrathin tip, followed by subsequent single-cell isolation (**Supplementary Figure 1**). *S. robusta* cells had naturally decreased in size by the time of the experiment and did not need experimental intervention.

RNA extraction protocol

Prior to the harvesting, cultures were inoculated into 182 cm2 vented cap cell culture flasks (VWR®) and cultured for approximately four days or until they reached the late exponential phase. In total 10 – 12 millions of cells were harvested per sample. Filters with cell biomass were preserved in 2ml Microcentrifuge tubes (Safe-Lock PCR clean, Eppendorf®), flash frozen in liquid nitrogen. The RNA from the samples was extracted with Qiagen RNeasy extraction kit and the manufacturer’s protocol was used with the following optimizations. Cells were scraped from the filter and resuspended in 600 µl of RLT buffer with 1 % of β-mercaptoethanol added. Silicon carbide beads were added to the tubes that were subsequently beaten in Mixer Mill MM 400 (Retsch GmbH) for 2 min 30 sec at 20 Hz. A DNase treatment was performed on-column according to the manufacturer’s instructions. Finally, the quality of the extracted RNA was assessed by a Thermo ScientificTM NanoDrop 2000 Spectrophotometer and the Agilent 2100 BioAnalyzer (using the Agilent RNA 6000 Pico Kit), and samples were stored at -80°C until further use.

Quality control of RNA-sequencing

Quality control was performed on all samples using FastQC version 0.11.2 (Babraham Bioinformatics, under GPL3 license). Adapter sequences were trimmed with Trimmomatic v0.36 (ILLUMINACLIPTruSeq3-PE.fa:2:30:10:2:true) (Bolger *et al.*, 2014).

Differential expression analysis of *C. closterium* size and mating type genes

A differential expression (DE) analysis was performed in Rstudio v3.36.0 for the size and mating type dataset of *C. closterium*. Isoform-level abundances were imported in R using tximport v1.22.0 (Soneson *et al.*, 2016) and aggregated to the gene level. Independent filtering was executed by keeping genes with at least one count per million (CPM) in at least three samples, thus retaining 19,201 *C. closterium* genes. Trimmed mean of M values (TMM) normalization was applied to correct differences in sequencing depth and RNA composition (Robinson & Oshlack, 2010). For each gene, negative binomial generalized linear comparing the contrasts model was fitted with EdgeR v3.36.0 (Robinson *et al.*, 2009). Next, likelihood ratio tests were performed across four different contrasts, and p-values were controlled at the 5 % FDR level using the Benjamini-Hochberg correction. In specific cases, genes were concurrently assigned to two of the four contrasts **(Supplementary Data 1**).

Differential expression analysis of *S. robusta* expression atlas and additional samples

A total of 198 RNA-seq samples from the *S. robusta* expression atlas and a 48-h diurnal cycle experiment (Bilcke *et al.*, 2021) were reprocessed by mapping to the *S. robusta* reference genome v1 with Salmon v1.3. After summarization on the gene level with tximport v1.22.0, the samples from control conditions (vegetative, untreated) were selected for differential expression analysis. After filtering and normalization, likelihood ratio tests were performed between 36 MT+ and 12 MT- samples against a logFC threshold of 6 using the GLMtreat command in edgeR. We tested against a log2 fold change threshold of six to select only strong mating type-dependent genes and avoid an overpowered analysis due to the high number of replicates. In order to visualize gene expression of selected genes in the other, non-control conditions of the gene expression atlas (biotic, environmental, diurnal, toxins, sexual reproduction) as well as samples from the six additional *S. robusta* strains discussed above, reads were imported into R with tximport v1.22.0 (Soneson *et al.*, 2016) and converted to counts per million (CPM) using the cpm command in edgeR.

Quality trimming, mapping and differential expression analysis of Sex-inducing pheromone (SIP) experiment

Transcriptomic datasets comprising 12 samples (6 control, 6 SIP filtrate treated) were first quality checked using FastQC version 0.11.9 (Babraham Bioinformatics, under GPL3 license), followed by adapter trimming using Trimmomatic v0.36 using the parameters given above (Bolger *et al.*, 2014). Trimmed paired-end reads were mapped to transcripts belonging to the gene annotation model v1.2 of the *C. closterium* reference genome (Audoor *et al.*, 2024) using Salmon v1.8 (Patro *et al.*, 2017) with following error correction modes enabled: --seqBias --gcBias. Isoform-level abundances were imported in R using tximport v1.30.0 (Soneson *et al.*, 2016) and aggregated to the gene level. Independent filtering was executed by keeping genes with at least one count per million (CPM) in at least three samples, thus retaining 18,575 *C. closterium* genes. Trimmed mean of M values (TMM) normalization was applied to correct differences in sequencing depth and RNA composition (Robinson & Oshlack, 2010). Generalized linear models were fitted for each gene and likelihood ratio tests were performed against a log2(fold change) cutoff of 1 using glmTreat in EdgeR (Robinson *et al.*, 2009). Differential expression was assessed across two contrasts: (1) between matched SIP-treated versus control samples after 3h of illumination, (2) between matched SIP-treated and control samples after 9h of illumination. The False Discovery Rate (FDR) was controlled at 5% using the Benjamini-Hochberg correction.

Functional annotation, phylostratigraphy and enrichment analysis

Taxonomic distribution, phylogenies and sequences of gene families can be explored through the PLAZA Diatoms 1.0 platform, e.g. https://bioinformatics.psb.ugent.be/plaza/versions/plaza_diatoms_01/gene_families/view/HOM02SEM009752. Transmembrane domains were predicted with Phobius ([phobius.sbc.su.se/](https://phobius.sbc.su.se/)), signal peptides were predicted with SignalP 6 (Teufel *et al.*, 2022) and other subcellular localization with HECTAR using the interactive Galaxy platform ((Gschloessl *et al.*, 2008), <https://webtools.sb-roscoff.fr/>). A nuclear localization signal was detected by LOCALIZER (Sperschneider *et al.*, 2017), and WoLF PSORT (Horton *et al.*, 2007). For each *C. closterium* gene, the phylostratigraphic age was determined by determining the phylogenetic scope of the gene family it belongs to. Five taxonomic bins were defined: a species distribution limited to *C. closterium*, pennate diatoms, diatoms (i.e. found in both pennate and centric diatoms) or extending to other, non-diatom, eukaryotes. Furthermore, a comparative transcriptomics analysis was performed by identifying gene families containing differentially expressed genes in response to mating type and/or size for both *C. closterium* and *S. robusta*. The enrichment of specific InterPro domains, gene families, and phylostrata among the differentially expressed genes of different size/mating type contrasts was determined using a hypergeometric test implemented in the enricher function from the ClusterProfiler package for R 4.10.0 (Wu *et al.*, 2021) using an FDR-adjusted p-value of 0.05 to define enrichment.

cDNA synthesis, RT-qPCR cycling conditions and statistical analysis of results

Total RNA was reverse transcribed using the iScript cDNA synthesis kit (Bio-RAD) following manufacturer’s instructions. Finally, an equivalent of 100 ng of reverse-transcribed RNA (cDNA) was used as a template in each qPCR reaction. Gene-specific primers (see **Supplementary  Table 2**) were designed towards the 3’ end of the target and control genes via the web-based Primer3Plus software (www.primer3plus.com), aiming for an amplicon length of 80–250 bp and a primer melting temperature Tm of 54–62°C. Samples were amplified in triplicate on the Lightcycler 480 platform (Roche) with Lightcycler 480 SYBR Green I Master mix (Roche Applied Science) in the presence of 5µM gene-specific primers. The cycling conditions were the following: 10 min pre-incubation at 95°C and 40 amplification cycles of 10 s at 95°C, 20 s at 58°C and 20 s at 72°C. After cycle 40, a melting curve analysis was carried out by heating from 65 to 97°C (0.5°C increments, 10 s dwelling time) to check for primer dimers and non-specific amplification. Relative expression levels were calculated using the delta-delta Ct quantification method in Excel. The stably expressed *histone* *h2b* (Histone H2B protein), *tubulin β* Tubulin β) and *tbp* (TATA binding protein) genes were used for normalization.

Hidden Markov model searches and phylogenetic analysis for MRM2 homologs

In order to execute phylogenetic analysis, three genes in the PLAZA MRM2 gene family with incorrect gene models were first manually curated using GenomeView (Abeel *et al.*, 2012). *Gene10933* and *ps00G37970* were extended because they were truncated and *ptri77580* was merged with *ptri77600*.  Protein sequences of these updated gene models are included as **Supplementary Data**. Annotation of protein domains in these updated gene models was performed with InterProScan (ebi.ac.uk/interpro/search/sequence, accessed on August 8^th^ 2023). An alignment of HOM02SEM009752 proteins including the curated sequences was created using MAFFT v7.453 (Katoh & Standley, 2013). After trimming in MEGA X to retain the conserved LRR domain (Kumar *et al.*, 2018),  a profile HMM was created for the MRM2 family using the hmmbuild command of HMMER (hmmer.org) which is included in **Supplementary Data** as a reference for the MRM2 family. A targeted search for MRM2 homologs in eukaryotes was performed using the hmmsearch command of HMMER, querying the MRM2 profile HMM against the proteome of 28 species, which include the 26 species included in the PLAZA Diatoms database (bioinformatics.psb.ugent.be/plaza/versions/plaza_diatoms_01/), as well as *C. closterium* (Audoor *et al.*, 2024) and *S. marinoi* (Pinder *et al*., in prep., with accession JATAAI000000000). Phylogenetic analysis was performed using the top-50 proteins that were selected based on the highest domain-specific E-value score. In brief, an alignment was created with MAFFT v 7.453 (Katoh & Standley, 2013), automatic trimming was performed with TrimAl v 1.4.1 (Capella-Gutiérrez *et al.*, 2009) with a gap threshold of 0.5, and a maximum-likelihood, midpoint rooted phylogenetic tree was generated with IQ-tree v 2.2.2.6 (Nguyen *et al.*, 2015), using 1000 ultrafast bootstrap repeats. The bootstrap consensus tree was midpoint rooted with the phangorn package for R v2.11.1 (Schliep, 2011), and visualized with ggtree v3.10.0 (Yu *et al.*, 2017). Protein structures were visualized with DrawProteins v3.18 (Brennan, 2018).
