## Supplementary Figures for "Molecular fingerprints of cell size sensing and mating type differentiation in pennate diatoms"

Supplementary Material

| **Species** | **Strain name** | **BCCM/DCG accession number** | **Place of origin / Parenthood of laboratory produced strains** | **Collector** | **Year of origin** | **Mating type (MT)** |
| --- | --- | --- | --- | --- | --- | --- |
| *C. closterium* | CA1.15 | DCG 0923 | Original strain, France, Baie de la Canche, Touquet-Paris-Plage | Willem Stock | 2014 | MT- |
| *C. closterium* | ZW2.20 | DCG 0922 | Original strain, Belgium, Zwin, Hoofdgeul | Willem Stock | 2014 | MT+ |
| *C. closterium* | NH3.13 | DCG 0685 | Original strain, Netherlands, Waddenzee, near entrance Amstelmeer | Willem Stock | 2014 | MT- |
| *C. closterium* | WS3.7 | DCG 0623 | Original strain, Netherlands, Wester-Schelde, Paulina Polder | Willem Stock | 2014 | MT+ |
| *C. closterium* | A5 | - | Cross of CA1.15 x ZW2.20 | Darja Belisova | 2018 | MT- |
| *C. closterium* | A6 | DCG 0980 | Cross of CA1.15 x ZW2.20 | Darja Belisova | 2018 | MT- |
| *C. closterium* | C2 | - | Cross of CA1.15 x ZW2.20 | Darja Belisova | 2018 | MT+ |
| *C. closterium* | MC3 | - | Cross of NH3.13 x WS3.7 | Darja Belisova | 2018 | MT- |
| *C. closterium* | MC4 | - | Cross of NH3.13 x WS3.7 | Darja Belisova | 2018 | MT+ |
| *C. closterium* | MC5 | - | Cross of NH3.13 x WS3.7 | Darja Belisova | 2018 | MT+ |
| *S. robusta* | 85 A | DCG 0105 | Progeny, Belgium, Oost-Vlaanderen, Gent, Ghent University Campus Sterre - S8, Krijgslaan, | Victor Chepurnov | 2006 | MT+ |
| *S. robusta* | VM 3-12 | DCG 0515 | Original strain, Netherlands, Veerse Meer | Sam De Decker | 2014 | MT- |
| *S. robusta* | Ponton 36 | DCG 0462 | Original strain, Netherlands, Veerse Meer | Peter Vanormelingen | 2012 | MT+ |
| *S. robusta* | VM 3-32 | DCG 0518 | Original strain, Netherlands, Veerse Meer | Sam De Decker | 2014 | MT- |
| *S. robusta* | 85A-1 | - | Cross of 85 A x VM 3-12 | Darja Belisova | 2018 | MT- |
| *S. robusta* | 85A-6 | - | Cross of 85 A x VM 3-12 | Darja Belisova | 2018 | MT- |
| *S. robusta* | 85A-9 | - | Cross of 85 A x VM 3-12 | Darja Belisova | 2018 | MT+ |
| *S. robusta* | P4 | - | Cross of Ponton 36 x VM 3-32 | Darja Belisova | 2018 | MT- |
| *S. robusta* | P5 | - | Cross of Ponton 36 x VM 3-32 | Darja Belisova | 2018 | MT+ |
| *S. robusta* | P7 | - | Cross of Ponton 36 x VM 3-32 | Darja Belisova | 2018 | MT+ |

**Supplementary Table 1: Information on diatom strains used in this study.** Place of origin and parenthood of laboratory maintained strains is indicated. All strains, produced as laboratory crosses, were isolated and maintained at the Protistology and Aquatic Ecology Lab in the S8 building at Campus Sterre, Krijgslaan 8, Gent. BCCM/DCG accession number was assigned to strains when included in Diatoms Collection by Belgian Coordinated Collections of Microorganism and enables a search of a particular strain in the BCCM/DCG database.

| **Gene** | **Primer sequence** | **Strand** | **Primer melting temperature** | **Amplicon length** |
| --- | --- | --- | --- | --- |
| Ccl_5938 (TUB-b) | ACGGACGCTACTTGACTTGC | forward | 57.6 °C | 167 bp |
|  | CTCAAGTCCCTTTGGTGGAAC | reverse | 55.8 °C |  |
| Ccl_2393 (H2B) | TGAACGCATTGCCACTGAAG | forward | 56.2 °C | 89 bp |
|  | AACGGACAGCGGTTTGAATC | reverse | 55.9 °C |  |
| Ccl_21857 (TBP) | ACAGCTCTCATTTTCGAGTGG | forward | 55 °C | 104 bp |
|  | ACGCGTTCGATGATGTAATGG | reverse | 55.5 °C |  |
| Ccl_4437 | CTGCTTCTCGTCCCAATCTC | forward | 55.3 °C | 170 bp |
|  | ATACGCAATTCCGAGGACAG | reverse | 54.8 °C |  |
| Ccl_5484 | CATGTGTGCCAATCTTGACC | forward | 54.3 °C | 161 bp |
|  | CATGGTTGCCATGATGAGAC | reverse | 53.9 °C |  |
| Ccl_13556 | GGTTTTCTGTGGCTGTGGAT | forward | 55.6 °C | 154 bp |
|  | ACATTTGTGATGTGGGCTGA | reverse | 54.8 °C |  |
| Ccl_14859 | ATCAAGTGCGAAGACGTTGG | forward | 58.9 °C | 139 bp |
|  | TGTGTTGTTCGATGGTTGCC | reverse | 59.3 °C |  |
| Ccl_14861 | TCAATGCCCACGATCGCATC | forward | 61.2 °C | 141 bp |
|  | CGACAACATCCTTCAGATCACG | reverse | 59.4 °C |  |
| Ccl_16260 | AAGGAAGTGGCAAGGGATCT | forward | 56.2 °C | 150 bp |
|  | GGGTGGTTACGGGACTTTTT | reverse | 55.3 °C |  |

**Supplementary Table 2: Primers used in the RT-qPCR experiment**. TUB-b: Tubulin β homolog, H2B: Histone H2B homolog, TBP: TATA binding protein homolog.

|  | large MT- A5 | large MT- A6 | large MT- MC3 | large MT+ C2 | large MT+ MC4 | large MT+ MC5 | small MT- A5 | small MT- A6 | small MT- MC3 | small MT+ C2 | small MT+ MC4 | small MT+ MC5 |
| --- | --- | --- | --- | --- | --- | --- | --- | --- | --- | --- | --- | --- |
| Ccl_14859 | 1.7 | 0.2 | 0.9 | 0.7 | 9.8 | 0.0 | 0.0 | 1.5 | 1.5 | 28.2 | 544.3 | 156.8 |
| Ccl_14861 | 0.0 | 0.0 | 0.0 | 0.0 | 0.9 | 0.0 | 1.7 | 2.1 | 0.0 | 19.4 | 474.6 | 98.3 |
| Ccl_14874 | 0.0 | 1.4 | 0.6 | 0.2 | 6.0 | 2.6 | 0.4 | 3.8 | 1.0 | 25.2 | 972.7 | 315.2 |

**Supplementary Table 3: expression in counts per million of three *C. closterium* genes** (rows). Each column represents a single replicate sample, indicating the cell size class, the mating type (MT) and genotype of each sample.

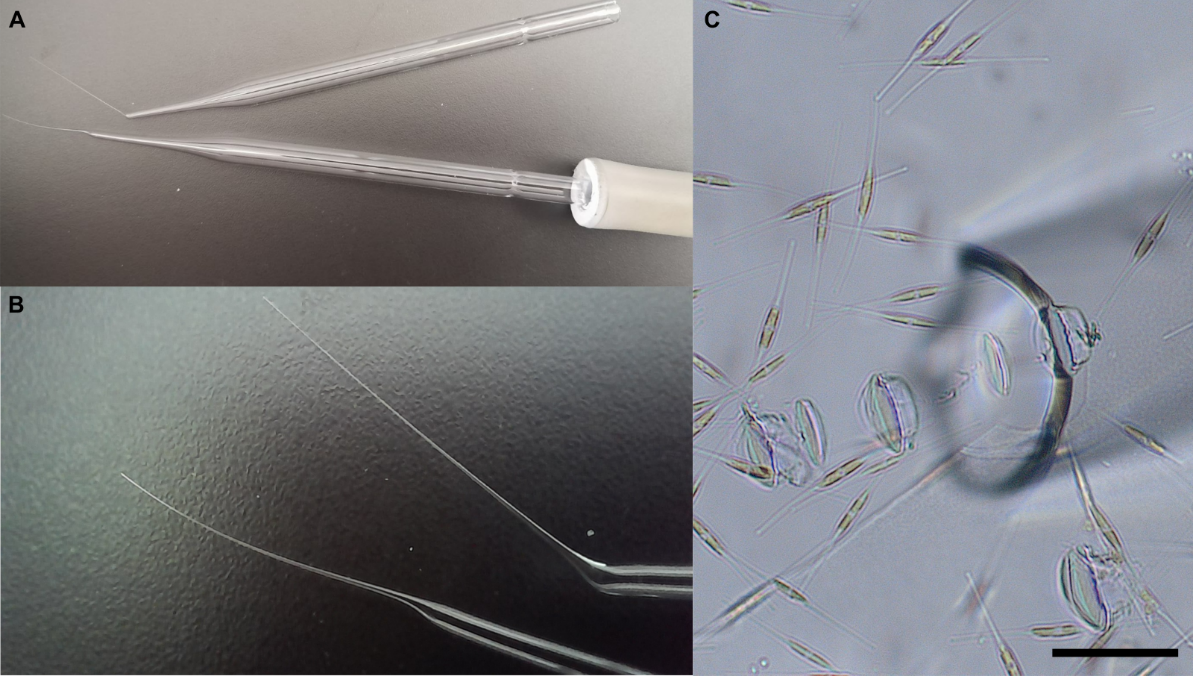

**Supplementary Figure 1: A**)**, B**) Glass Pasteur pipettes with a narrowed tip, prepared in the laboratory using Bunsen Burner. **C**) Light microscopy image showing *C. closterium* cells together with the sharp and open tip of the Pasteur pipette that was used to cut off the cell apices. Individual cells for which apices were successfully cut were subsequently isolated and used to start new clonal cultures. Scale bar 50 µm.

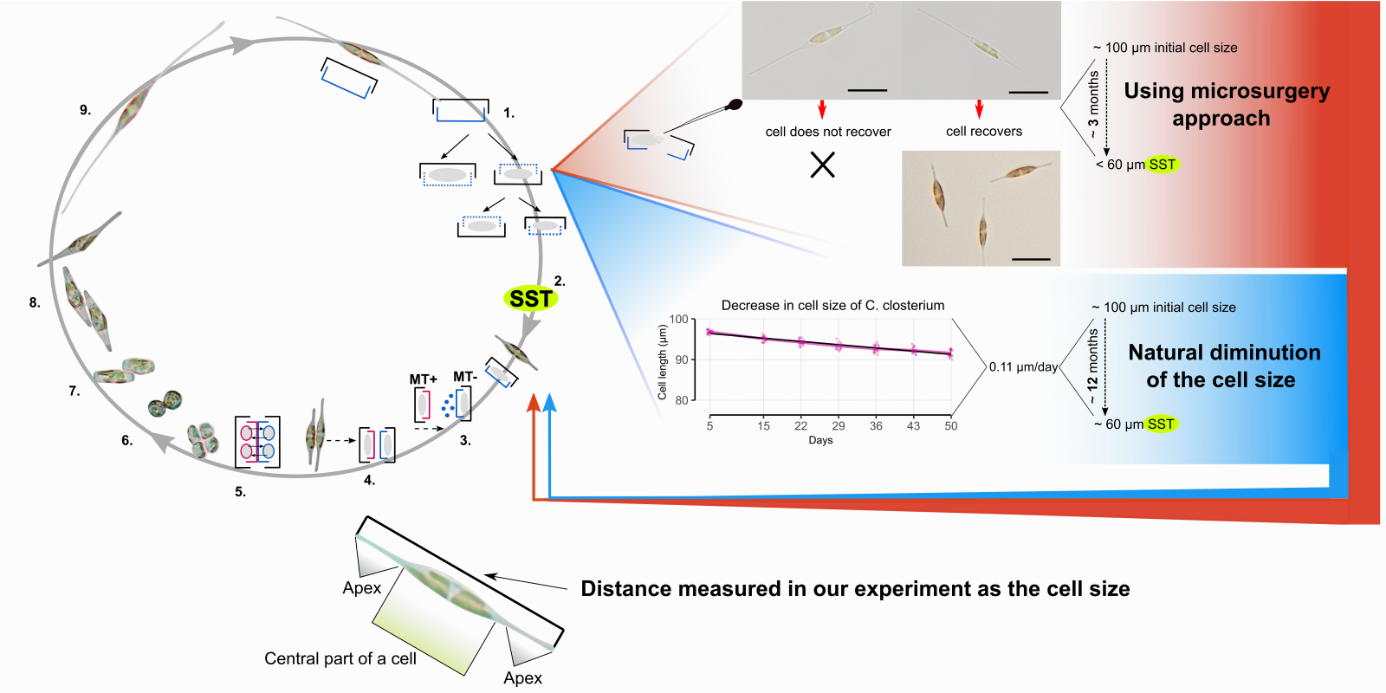

 **Supplementary Figure 2: Life cycle of *C. closterium* with two alternative ways of how cells decrease in size schematically described.** A natural decrease in the cell size (blue colour) can be estimated from a pilot experiment where we regularly measured the cell size of *C. closterium* strain B7  over 50 days. Cells decrease in cell size linearly by 0.11 µm/day, meaning the time necessary to reach SST is approximately one year. This time can be significantly shortened when using microsurgical removal of cell apices (red colour) (scale bar 20 µm). The distance between the tips of the apices was chosen as a parameter to measure the total cell size in our experiments.

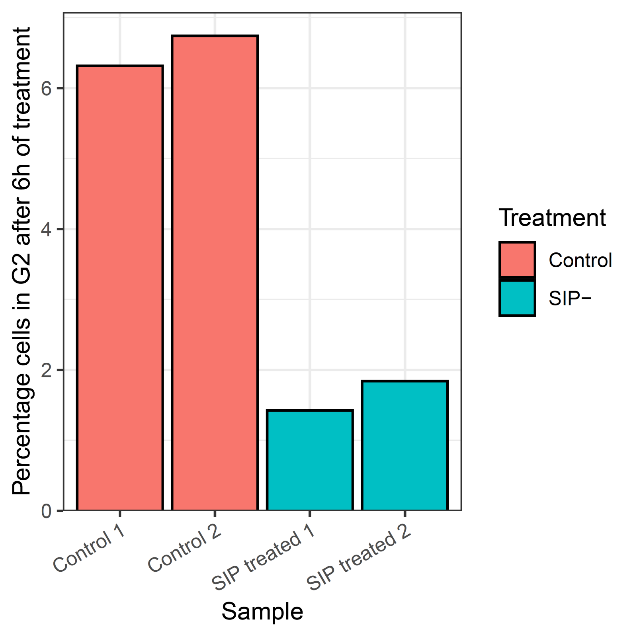

**Supplementary Figure 3: bar plot showing the results of an activity assay for SIP- containing medium, where a MT+ culture (strain ZW2.20) was treated with SIP- containing medium for 6h.** Different bars represent individual samples (n = 2). The y-axis shows the proportion of cells that is in the G2/M phase of the cell cycle, as determined through flow cytometry. Cultures that were treated with SIP- containing medium are coloured in cyan.

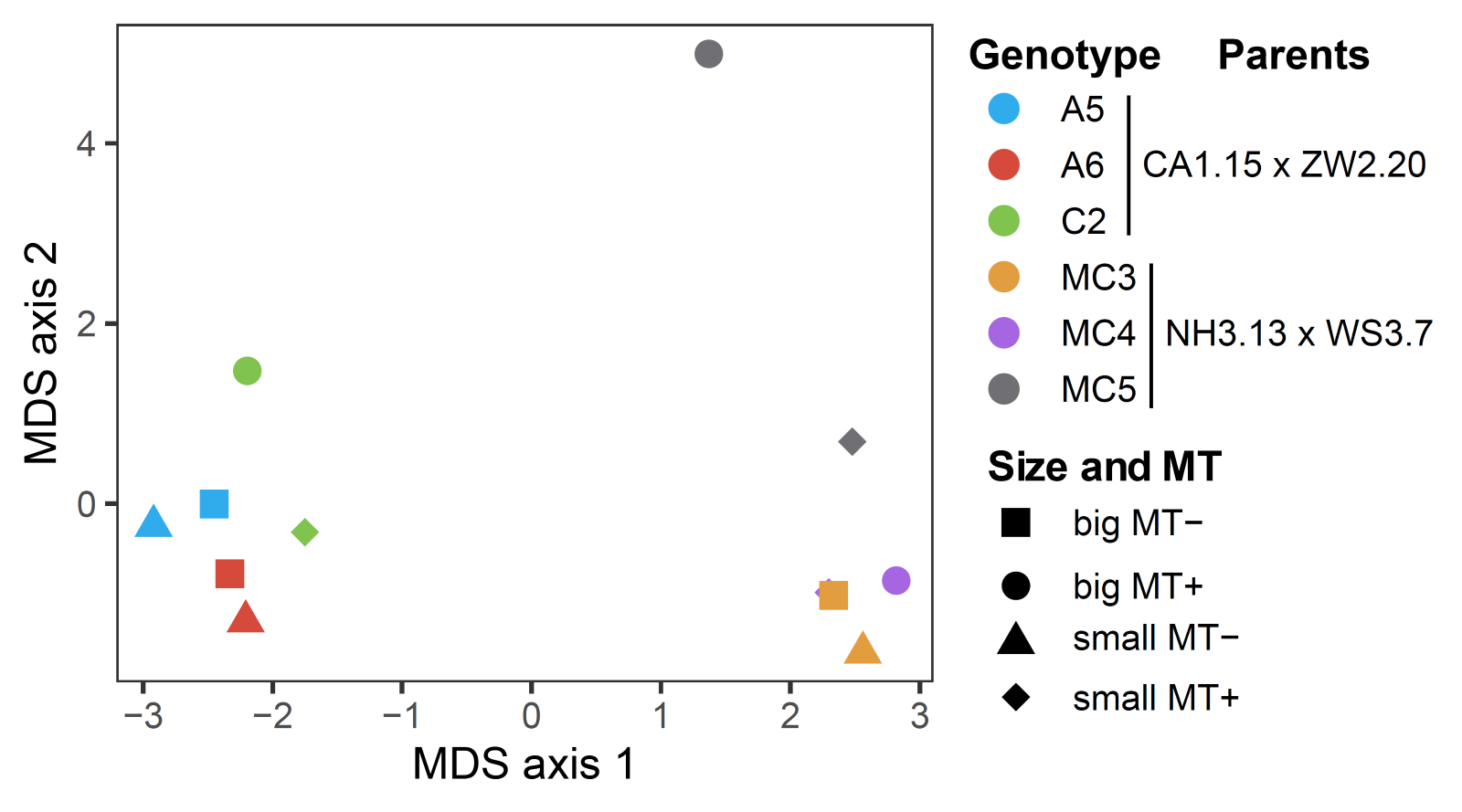

**Supplementary Figure 4: multidimensional scaling (MDS) plot for gene expression of 12 RNA-seq samples in *C. closterium*.** Shapes indicate the different size classes and mating types (MT), while colors indicate the different genotypes used in this RNA-seq experiment, with the parental strains (genetic lineage) indicated in the legend. The distance between samples resembles the log2 fold changes of the top-500 genes.

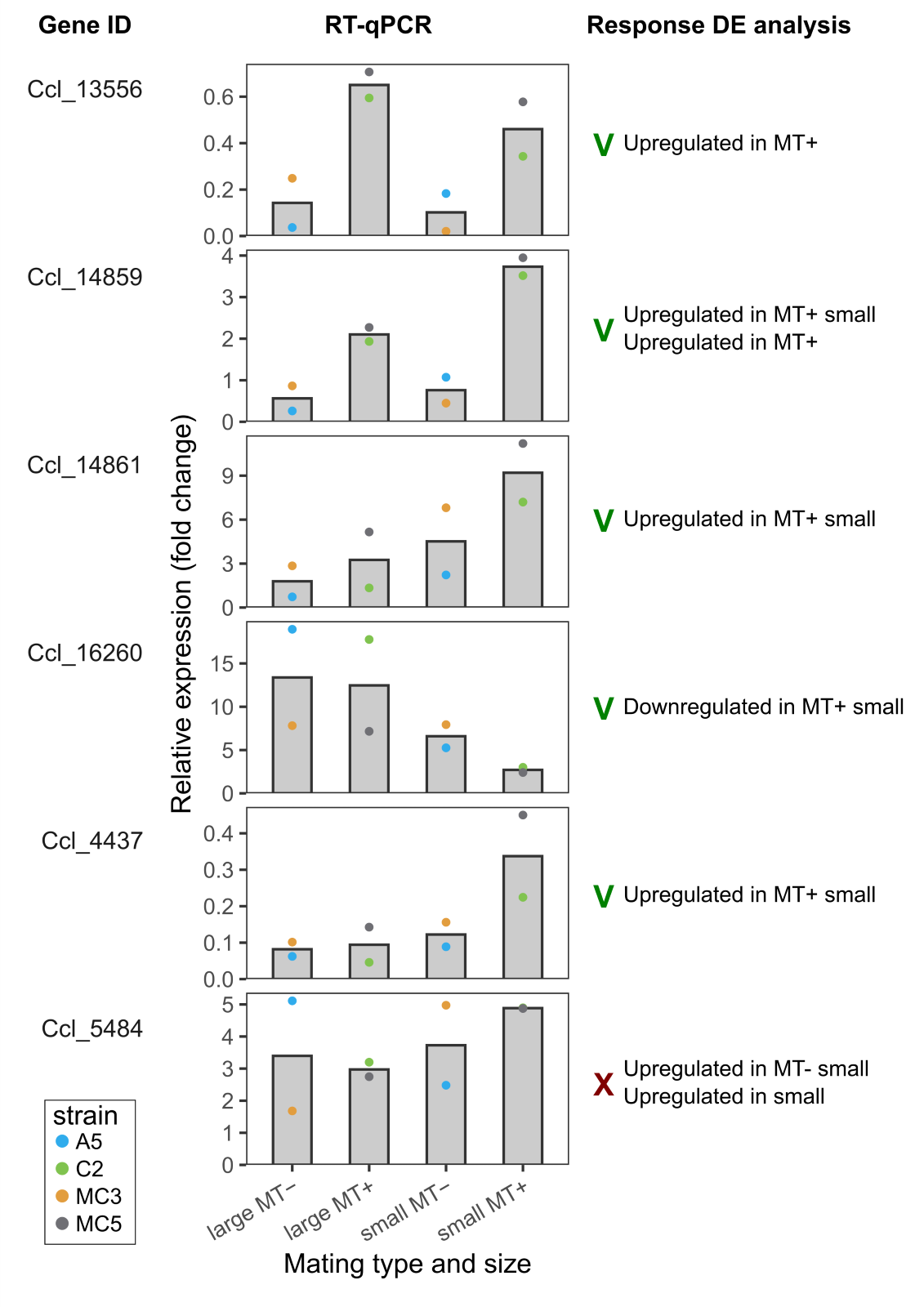

**Supplementary Figure 5: validation of the expression of six *Cylindrotheca closterium* genes using reverse transcriptase quantitative PCR (RT-qPCR).** Relative expression is shown for cultures in four different size and mating type classes: large MT-, large MT+, small MT- and small MT+. Bars show the average expression, while points indicate the expression of individual replicates (coloured by strain, n = 2).

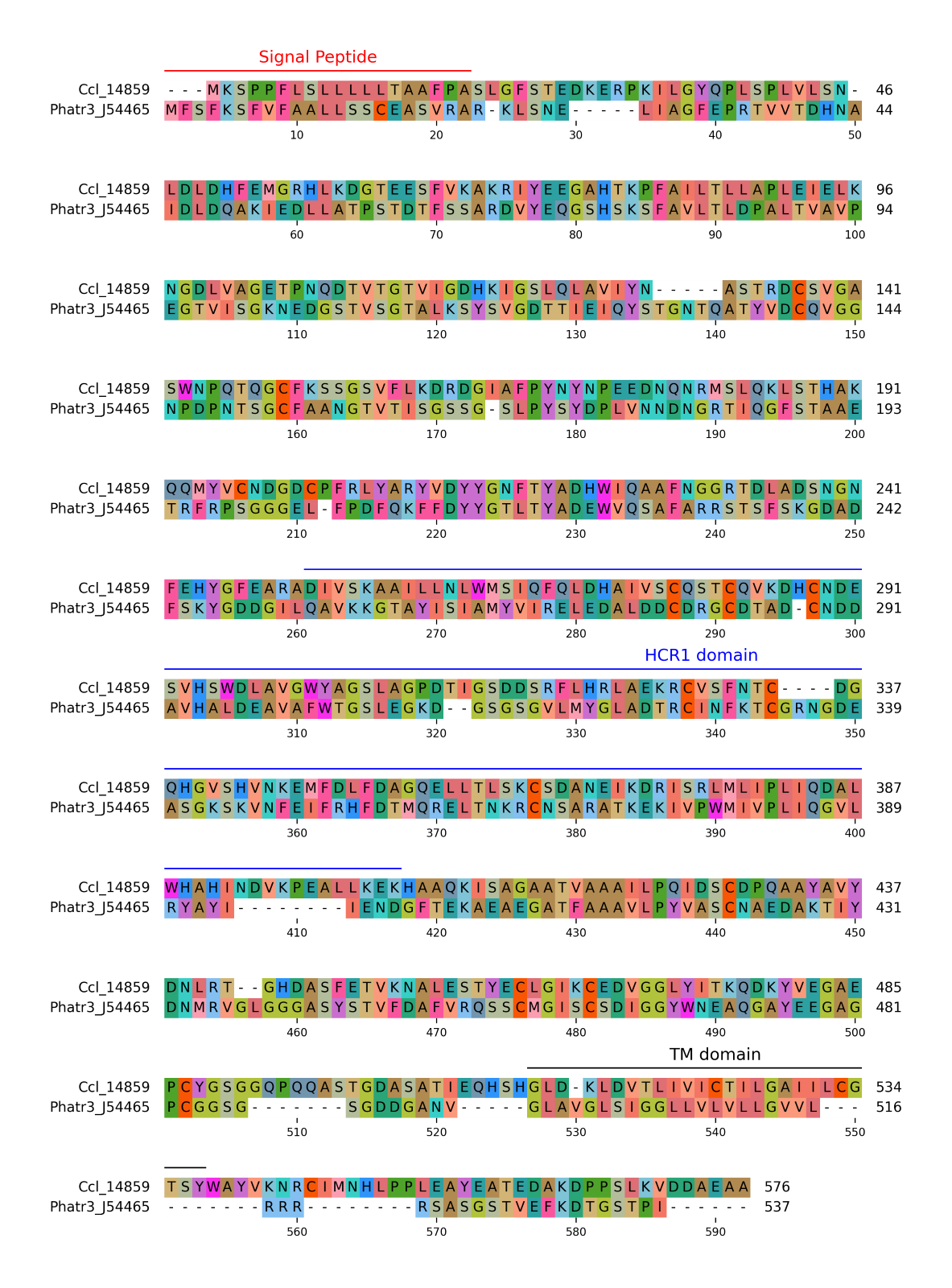

**Supplementary Figure 6: multiple sequence alignment of Ccl_14859 and *Phaeodactylum tricornutum* phytotransferrin Phatr3_J54465.** Alignment created with MAFFT 7.453 and vizualized with PyMsaViz 0.4.2 [1]. Protein domain prediction was performed with deepTMHMM 1.0 and InterProScan v 5.69-101.0.

[1] Shimoyama, Yuki. pyMSAviz: MSA visualization python package for sequence analysis. https://github.com/moshi4/pyMSAviz

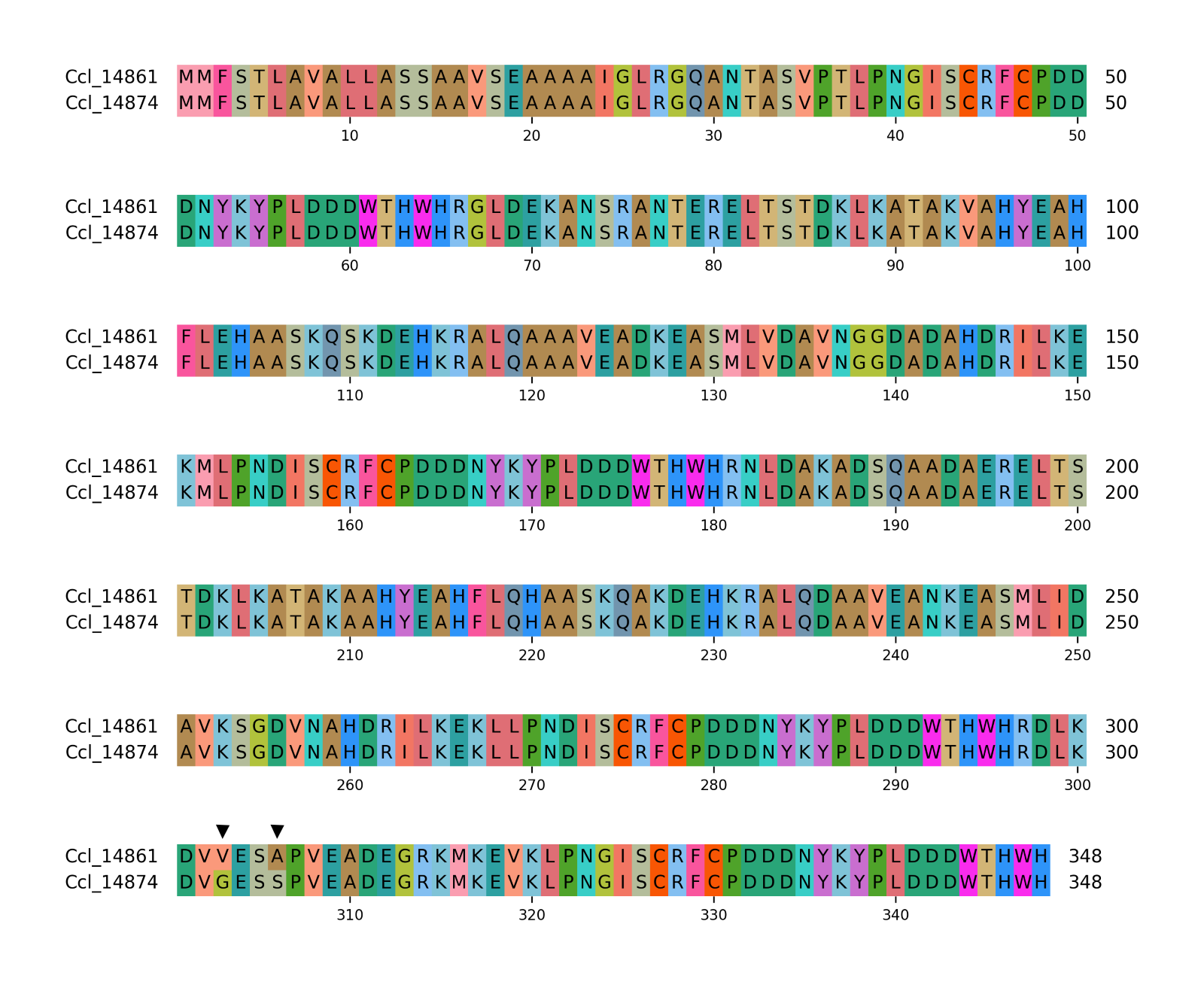

**Supplementary Figure 7: multiple sequence alignment of Ccl_14861 and Ccl_14874.** Alignment created with MAFFT 7.453 and vizualized with PyMsaViz 0.4.2 [1]. The different amino acids at positions 303 and 306 are indicated by black arrowheads.

[1] Shimoyama, Yuki. pyMSAviz: MSA visualization python package for sequence analysis. https://github.com/moshi4/pyMSAviz

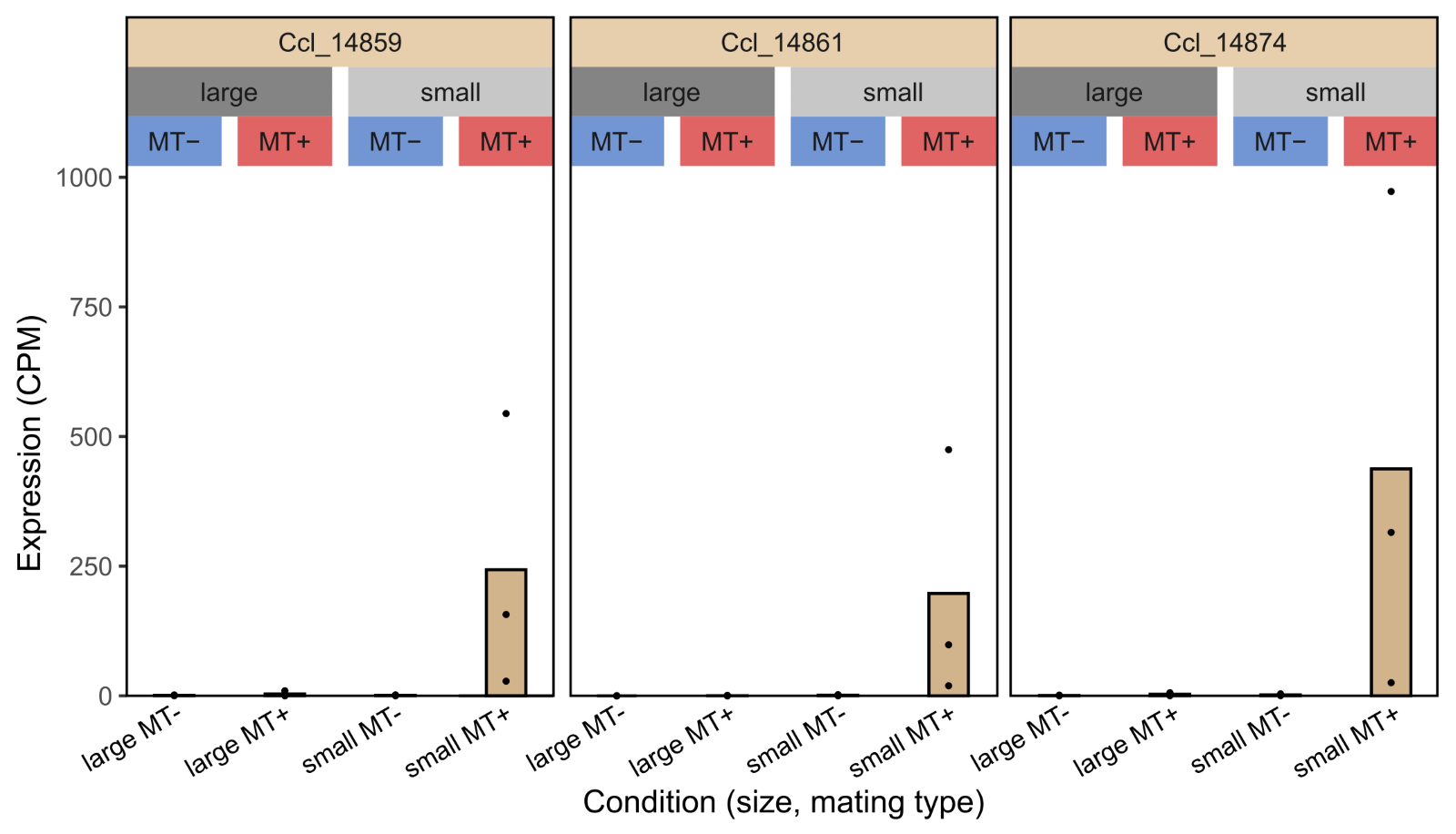

**Supplementary Figure 8: expression in counts per million (CPM) of three *Cylindrotheca closterium* genes in different mating types (MT) and across cell sizes: large cells larger than the sexual size threshold (SST) and small cells below the SST.** Bars show the average expression, while points show individual replicates (genotypes).

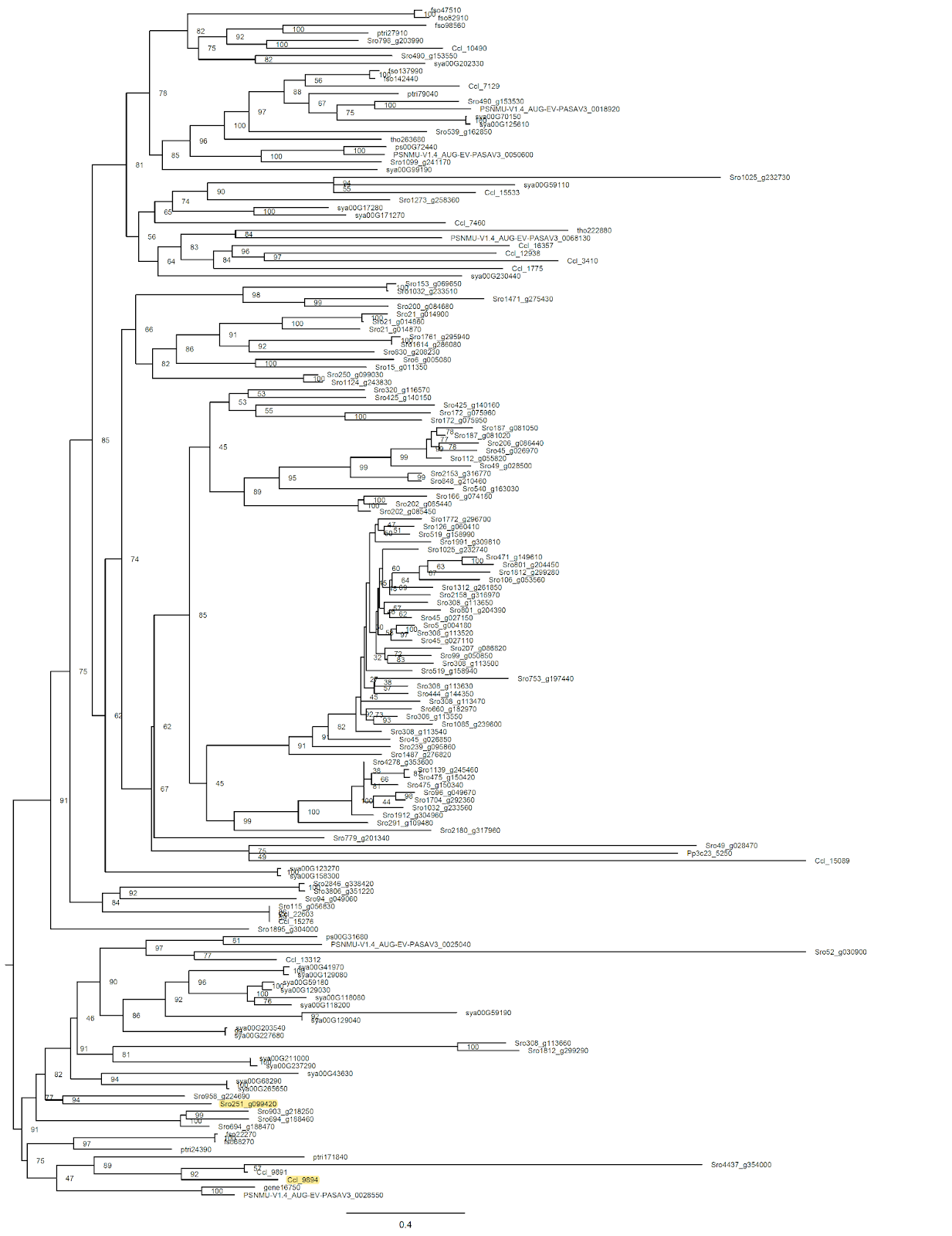

**Supplementary Figure 9: midpoint rooted bootstrap-consensus phylogenetic tree of the gene family HOM02SEM000188.** Bootstrap support is shown as node labels. Two genes with MT+ biased expression are highlighted in yellow.

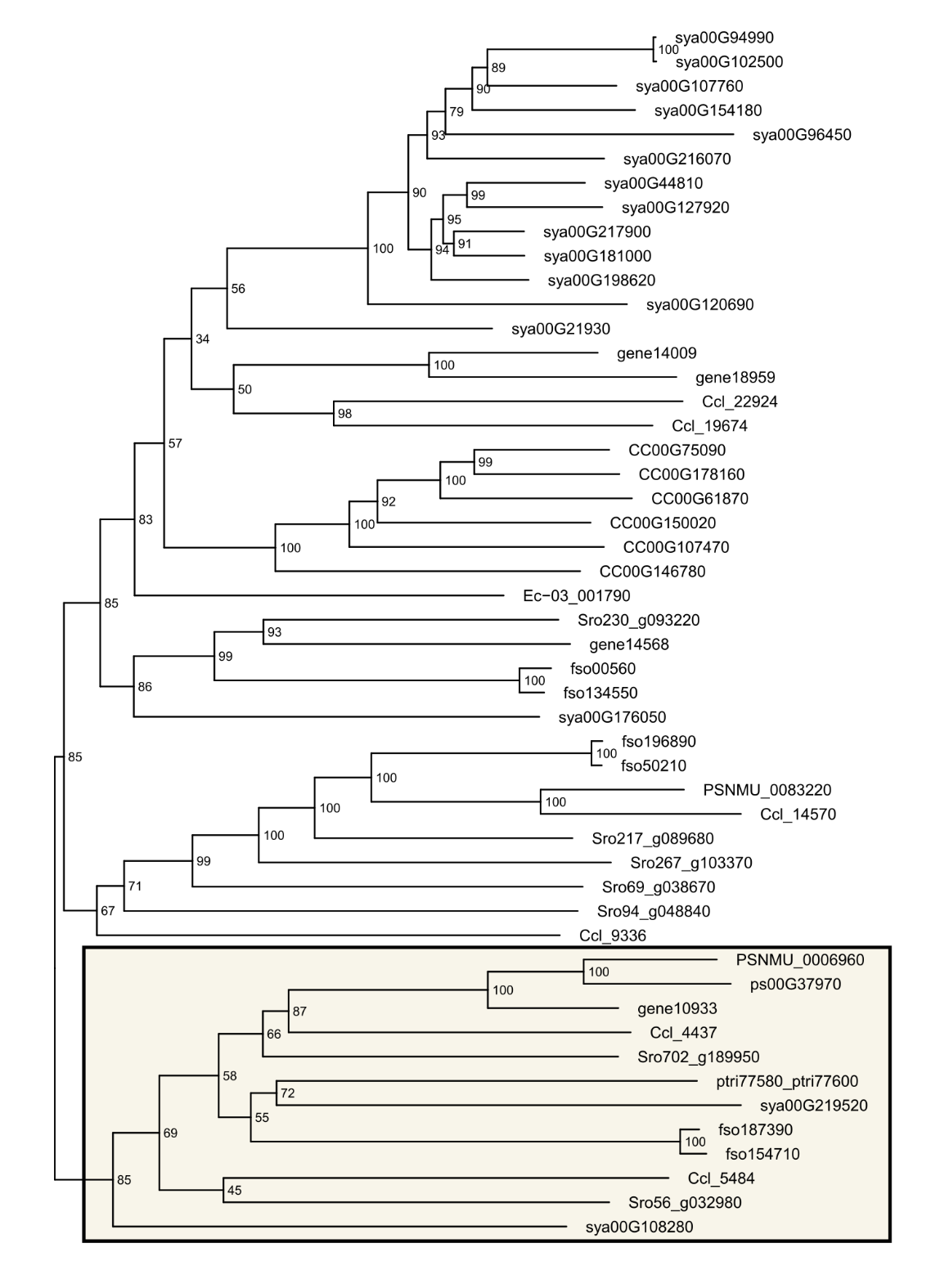

**Supplementary Figure 10: midpoint rooted bootstrap-consensus phylogenetic tree of top-50 proteins with the highest similarity to the MRM2 Leucine-rich repeat domain.** Node labels show bootstrap values. The box indicates the selected clade with MRM2 homologs.

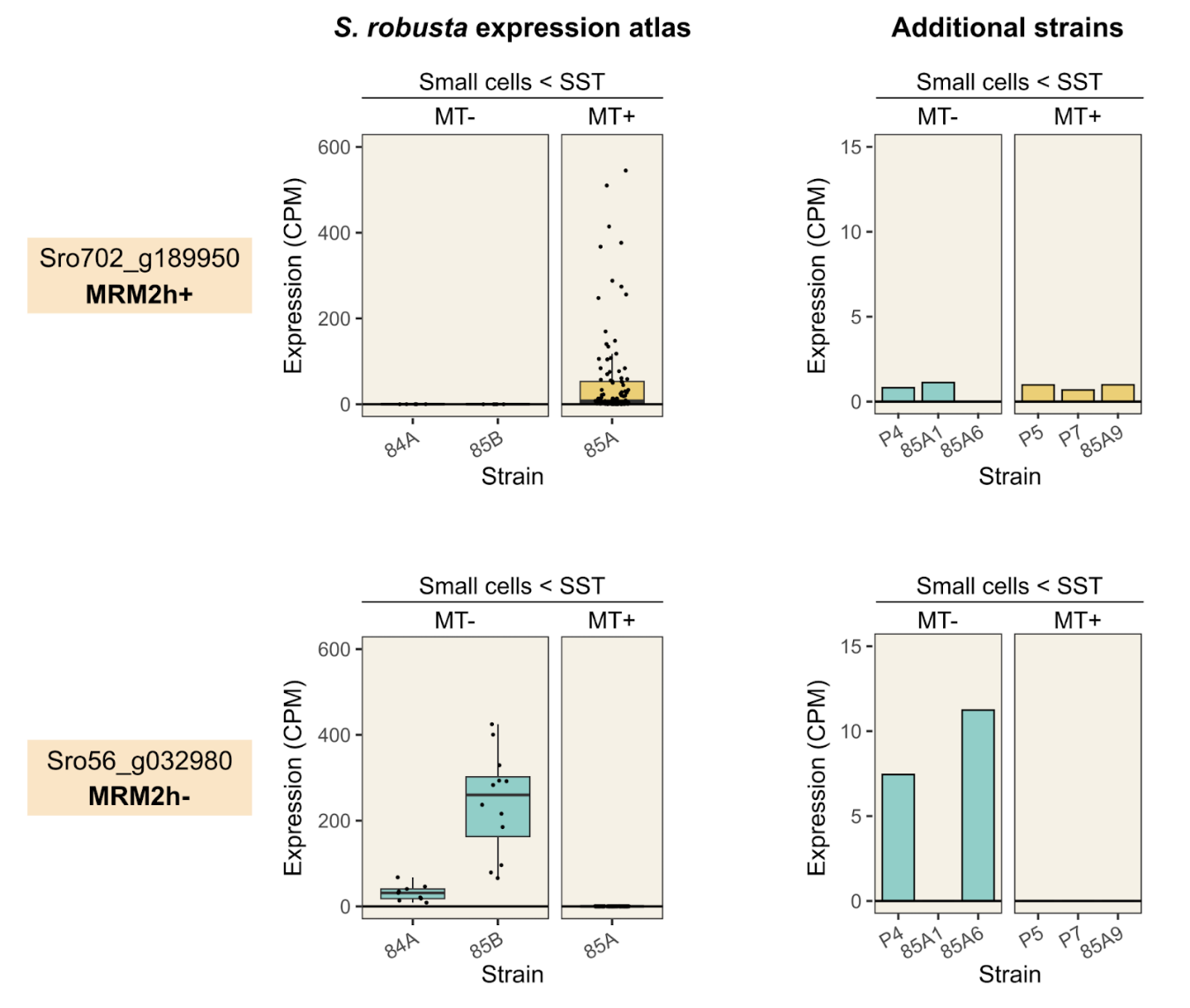

**Supplementary Figure 11: expression of MRM2 homologs in different genotypes (strains) of *Seminavis robusta***. Expression is shown as counts per million (CPM).
